# Changes of the biophysical properties of voltage-gated Na^+^ currents during maturation of the sodium-taste cells in rat fungiform papillae

**DOI:** 10.1101/2021.05.06.442879

**Authors:** Albertino Bigiani, Roberto Tirindelli, Lorenzo Bigiani

## Abstract

Taste cells are a heterogeneous population of sensory receptors that undergoes a continuous turnover. Different chemo-sensitive cell lines rely on action potentials to release the neurotransmitter onto nerve endings. The electrical excitability is due to the presence of a tetrodotoxin-sensitive, voltage-gated sodium current (*I_Na_*) similar to that found in neurons. Since the biophysical properties of *I_Na_* change during neuronal development, we wondered whether the same also occurred in taste cells. Here, we used the patch-clamp recording technique to study *I_Na_* in sodium sensing cells of rat fungiform papillae. We identified these cells by exploiting the known blocking effect of amiloride on ENaC, the sodium (salt) receptor. Then, based on the amplitude of *I_Na_* and a morphological analysis, we subdivided sodium cells into two broad developmental groups, namely immature and mature cells. We found that: the voltage dependence of activation and inactivation changed in the transition from immature to mature state (depolarizing shift); the membrane capacitance significantly decreased in mature cells, enhancing the density of *I_Na_*; a persistent sodium current, absent in immature cells, appeared in mature cells. mRNA expression analysis of the α-subunits of voltage-gated sodium channels in fungiform taste buds supported the electrophysiological data. As a whole, our findings provide evidence for a noticeable change in membrane excitability during development, which is consistent with the key role played by electrical signaling in the release of neurotransmitter by mature sodium cells.

**Key Points Summary:**

- Taste cells are sensory receptors that undergo continuous turnover while they detect food chemicals and communicate with afferent nerve fibers.
- The voltage-gated sodium current (*I_Na_*) is a key ion current for generating action potentials in fully differentiated and chemo-sensitive taste cells, which use electrical signaling to release neurotransmitters.
- Here we report that in rat taste cells involved in salt detection, the properties of *I_Na_*, such as voltage dependence of activation and inactivation, undergo significant changes during the transition from immature to mature state.
- Our results help understand how taste cells gain electrical excitability during turnover, a property critical to operate as chemical detectors that relay sensory information to nerve fibers.

## Introduction

Taste cells, the peripheral detectors of food chemicals, are unique among sensory receptors because they possess both neuronal-like and epithelial-like properties. Chemo-sensitive taste cells express several voltage-gated ion currents similar to those identified in neurons, including a tetrodotoxin (TTX)-sensitive sodium current, *I_Na_*, which generates action potentials (Béhé *et al*., 1990). These electrical signals are necessary for releasing the neurotransmitter onto nerve fiber endings (Taruno *et al*., 2013; Nomura *et al*., 2020). Indeed, firing capability is an electrophysiological hallmark of mature, chemo-sensitive taste cells (Béhé *et al*., 1990; Avenet & Lindemann, 1991; Gilbertson *et al*., 1992; Chen *et al*., 1996; Furue & Yoshii, 1997; Bigiani *et al*., 2002; Noguchi *et al*., 2003; Yoshida *et al*., 2006; Yoshida *et al*., 2009a; Yoshida *et al*., 2009b; Niki *et al*., 2011; Nomura *et al*., 2020; Ohtubo, 2021). Although taste cells are exquisitely specialized in terms of membrane excitability, however, their lifespan only lasts few weeks as they turnover like a regular epithelium (Barlow, 2015). Mature cells arise from post-mitotic ovoid cells at the base of the taste buds, the end organs of taste. These immature cells differentiate to produce elongated spindle-shaped cells that eventually reach the taste pore, where interaction with chemical stimuli occurs (Feng *et al*., 2014; Barlow, 2015; Yang *et al*., 2020). Therefore, taste buds in adult animals contain intermingled elements at different stages of growth.

The biophysical properties of *I_Na_* change significantly during the development of excitable cells. Typically, the maturation of this current consists of a gradual increase in amplitude along with changes in the voltage dependence of the activation and inactivation processes, which affect membrane excitability in fully differentiated cells (Cummins *et al*., 1994; Costa, 1996; Gao & Ziskind-Conhaim, 1998; Fry, 2006; Donnelly, 2011; Yu *et al*., 2011; Tadros *et al*., 2015; Calvigioni *et al*., 2017). Apart from a study on the amphibian *Necturus maculosus* (Mackay-Sim *et al*., 1996), it is mostly ignored whether and how *I_Na_* changes in mammalian taste cells during their maturation. The main reason is that chemo-sensitive taste cells are heterogeneous and the ability to fire action potentials is shared by separate cell populations with distinct molecular and functional properties (Chaudhari & Roper, 2010; Bigiani & Prandi, 2011). Thus, it is difficult to trace a given cell line during turnover in living preparations.

To circumvent this problem, we exploited electrophysiological markers to identify defined groups of taste cells. First, we selected cells with *I_Na_* and expressing the epithelial sodium channel (ENaC), a sodium receptor involved in detecting Na^+^ in rodents (Chandrashekar *et al*., 2010; Nomura *et al*., 2020). These sodium salt sensing cells (*sodium cells*) (Nomura *et al*., 2020) can be functionally identified by recording the so-called “response to amiloride” (*I_Am_*), a reduction in the stationary ENaC-mediated current when this inhibitor is applied to the cell membrane (Bigiani, 2016). Then, amongst sodium cells, we identified two subpopulations that largely differ for their *I_Na_* and that we defined as mature and immature elements. Thus, we were able to use patch-clamp recording techniques to study the biophysical properties of *I_Na_* in sodium cells from rat fungiform papillae at different developmental stages. In particular, we aimed at establishing whether during their maturation the voltage dependence of activation and inactivation changed. These processes are key factors in determining the membrane excitability, that is, the ability of cells to generate action potentials (Angelino & Brenner, 2007).

## Materials and methods

### Ethical approval

Experiments were performed in compliance with the Italian law on animal care No. 116/1992, and in accordance with the European Community Council Directive (EEC/609/86). All efforts were made to reduce both animal suffering and the number of animals used. This study was approved by the institutional review board, ‘Organismo Preposto al Benessere degli Animali (OPBA)’.

### Animals

Sprague-Dawley rats (male and female, 60-90 days of age) were used in this study. Animals were obtained from Envigo (San Pietro al Natisone, Italy) and were housed two per cage on a 12-h light/dark cycle in climate-controlled conditions with *ad libitum* access to water and food. We used the Sprague-Dawley rat as an animal model because the ENaC-mediated salt component of the taste nerve response is much greater than in other rodent species, including mice (Halpern, 1998), and this is associated with the presence of large amiloride-sensitive sodium currents in fungiform taste cells (Bigiani, 2015, 2017). To isolate taste tissue, animals were deeply anesthetized by inhalation of isoflurane (Merial Italia, Milan, Italy), and then euthanized by cervical dislocation.

### Isolation of rat fungiform taste buds

Electrophysiological experiments were performed on taste cells in isolated taste buds. The procedure for obtaining single taste buds from fungiform papillae closely followed published protocols, *e.g.* (Béhé *et al*., 1990; Doolin & Gilbertson, 1996; Kossel *et al*., 1997). Taste buds were plated on the bottom of a chamber consisting of a standard glass slide onto which a silicon ring (thickness, 1–2 mm; internal diameter,15 mm) was pressed. We used glass slide pre-coated with poly-lysine (Fisher Scientific, Milan, Italy) to improve adherence of isolated taste buds to the bottom of the chamber. The chamber was placed onto the stage of an upright Olympus microscope (model BHWI) equipped with a 40x water-immersion objective, and taste buds were viewed with Nomarski optics. During the experiments, isolated taste buds were continuously superfused with Tyrode solution (containing in mM: 140 NaCl, 5 KCl, 2 CaCl_2_, 1 MgCl_2_, 10 glucose, 10 sodium pyruvate, and 10 HEPES; pH 7.4) by means of a gravity-driven system.

### Patch-clamp recording

Standard whole-cell patch-clamp recordings were made from taste cells as previously described, *e.g.* (Béhé *et al*., 1990; Doolin & Gilbertson, 1996; Kossel *et al*., 1997), at 22-25 °C. Signals were filtered at 5 kHz with the patch-clamp amplifier (Axopatch 1-D, Molecular Devices, Sunnyvale, CA) and sampling rate was adjusted according to the kinetics of membrane current: 100 kHz for fast currents (capacitive transients and *I_Na_*), 100 Hz for slow currents (amiloride responses). Data were acquired and analyzed with the pCLAMP9 software (Molecular Devices, Sunnyvale, CA). Recording pipettes were made from soda lime glass capillaries (Globe Scientific, Paramus, NJ) on a two-stage vertical puller (PP-830, Narishige, Tokyo, Japan). We used pipettes with resistances between 2.1 and 2.7 MΩ when filled with a standard pipette solution containing (in mM) 120 CsCl, 1 CaCl_2_, 2 MgCl_2_, 10 HEPES, 11 EGTA, and 2 ATPNa_2_, pH 7.2 adjusted with CsOH. Offset potentials were zeroed before seal formation. The liquid junction potential of ∼4 mV measured between pipette solution and Tyrode (bath) solution was not corrected. Patch pipette was positioned on taste cells by means of a water hydraulic micromanipulator (MHW-3, Narishige, Tokyo, Japan).

In cell-attached configuration, applying a voltage step produced a fast capacitive transient mainly due to the stray component resulting from immersion of the patch pipette in the bath solution. This capacitive current was reduced by coating the pipette with plastic and by electronic compensation before breaking the membrane patch underneath the pipette tip. In whole-cell configuration, cell capacitance (*C_M_*) was evaluated from the integral of the capacitive transient evoked by a 20-mV voltage step from a holding potential of −80 mV (Bigiani & Roper, 1995). Since functional ENaCs are responsible for a leakage current (Bigiani, 2016) that could affect *C_M_* measurements (Lindau & Neher, 1988), the capacitive transients were evoked in the presence of 1 μM amiloride, which blocks more than 90% of ENaC-mediated currents (Bigiani, 2016).

### Identification of taste cells expressing functional ENaC

The presence of functional ENaC in taste cells was monitored by studying the effect of bath-applied amiloride on the leakage current recorded at a holding potential of −80 mV (Bigiani, 2016). To ensure specificity, we used an amiloride concentration of 1 µM, which is above the inhibition constant for ENaC in these cells and below the dose range affecting other membrane transporters (Lindemann, 1996). We included in the analysis taste cells that exhibited an amiloride response ≥ 20 pA, unless otherwise indicated.

### Voltage-gated sodium currents

To isolate *I_Na_* in ENaC-expressing taste cells, voltage-gated K^+^ currents were blocked by using a Cs^+^-based pipette solution (Bigiani, 2017) in the presence of 1 µM amiloride, which was added to all bathing solutions (Bigiani, 2016). Blocking ENaC was also indispensable to avoid changes in the intracellular Na^+^ concentration, which could affect the Nernst equilibrium potential for sodium (*E_Na_*) and therefore *I_Na_* (Bigiani, 2016). After obtaining whole-cell configuration, we waited a few minutes for the equilibration between the pipette solution and taste cell cytosol. Recording started when sodium currents became stable (Costa, 1996).

The access resistance (*R_access_*) of the patch pipette is the main determinant of the series resistance that influences the behavior of voltage-gated currents (Armstrong & Gilly, 1992; Sontheimer & Ransom, 2002). Problems in clamping the membrane potential could easily arise when trying to voltage-clamp very large and fast-rising sodium currents (Armstrong & Gilly, 1992). In order to reduce *R_access_* we used patch electrodes with as small resistance as possible (Armstrong & Gilly, 1992; Sontheimer & Ransom, 2002). Thus, we were able to obtain stable whole-cell recordings with electrode resistances in the 2.1–2.7 MΩ range. To determine if the clamping conditions were appropriate, we followed the qualitative criteria based on the appearance of current waveforms as previously described, *e.g.*, (Armstrong & Gilly, 1992; Cummins *et al*., 1994; Costa, 1996; Tadros *et al*., 2015). Specifically, current amplitude had to increase gradually and traces had to rise smoothly upon application of voltage commands. Records unfulfilling these criteria were not included in the analysis.

The activation of voltage-gated sodium channels was studied by determining the conductance-voltage (G-V) relationship for *I_Na_*. This was obtained by evaluating, for each voltage step, *G_Na_ = I / (V - E_Na_)*, with *E_Na_* being the Nernst equilibrium potential for Na^+^ estimated in our experimental conditions, and then by fitting data to the Boltzmann function (Standen *et al*., 1994; Sontheimer & Ransom, 2002):

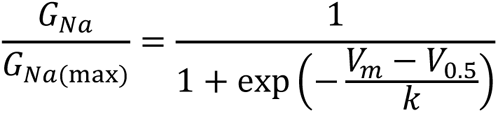

where: *G_Na_/G_Na(max)_* is the sodium conductance (*G_Na_*) normalized to its maximal value (*G_Na(max)_*), *V_m_* is the membrane potential at which the conductance was measured, *V*_0.5_ is the membrane potential at which conductance is 50 % of the maximal value, and *k* is the curve slope.

The voltage dependence of steady-state inactivation was studied following standard two-pulse procedure (Standen *et al*., 1994; Sontheimer & Ransom, 2002). Briefly, a first 500-ms pulse (prepulse) of potentials varying between - 120 and −20 mV was applied to the cell to develop inactivation, while a second pulse at −20 mV (test pulse) was used to measure the fraction of current that remained available for activation. The steady-state inactivation curves were obtained by fitting the data either to a single Boltzmann equation:

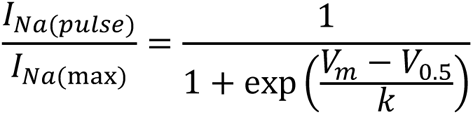

or to a double Boltzmann equation (Rush *et al*., 1998; Chen *et al*., 2000; Cummins *et al*., 2002):

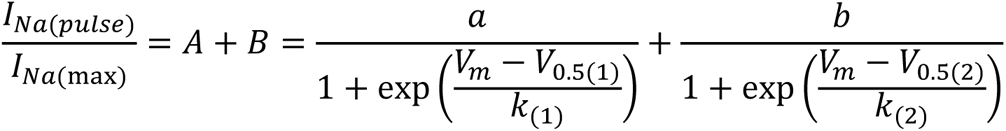

where: *I_Na(pulse)_*/*I_Na(max)_* is the sodium current elicited during a test pulse (*I_Na(pulse)_*) and normalized to the maximal current obtained with a prepulse potential of - 120 mV (*I_Na(max)_)*; *V_m_* is the voltage at which the membrane was held for 500 ms before the test pulse (prepulse potential). For single Boltzmann relationship, *V*_0.5_ is the membrane potential at which the current is 50 % inactivated, and *k* is the slope factor. For double Boltzmann relationship, subscripts 1 and 2 indicate first and second *V*_0.5_ and *k*, whereas *a* and *b* are fitting parameters that weight the influence of each single contribution (component “*A*” and component “*B*”) to the overall process, being *a* + *b* = 1. Data were considered as fitting to a double Boltzmann equation if the value of both *a* and *b* was > 0.05.

### Linear transformations

We used linearization analysis of data points to better visualize the presence of multiple components underlying a given process (Morton *et al*., 1995). The linearization of capacitive transients was obtained by plotting the logarithmic values of the falling phase of capacitive currents versus time. Plot was then analyzed by simple linear regression (Bigiani & Roper, 1995). The linearization of steady-state inactivation data was obtained by plotting the following parameter derived from the Boltzmann relationship versus the prepulse potential (*V_m_*):

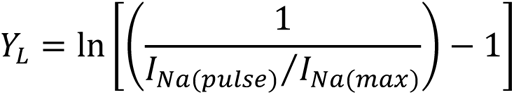

Straight lines were then fitted to the linear portions of the transformed data (Morton *et al*., 1995).

### Chemicals

All drugs were from Sigma-Aldrich (Milan, Italy). Amiloride was dissolved in dimethyl sulfoxide at a concentration of 0.5 M, and then diluted to its final concentration of 1 µM. Solutions were delivered through the gravity-driven flow perfusion system, which assured a solution exchange in less than 1 minute.

### RNA extraction and Reverse transcription polymerase chain reaction (RT-PCR)

To evaluate the expression of mRNA for sodium channel α-subunits in fungiform papillae, taste buds from three animals were isolated as above described and pooled together. Total RNA was extracted using TRIzol Reagent (Thermo Fisher Scientific) according to manufacturer’s instructions. RNA was then extensively treated with DNase I (Promega, Milan, Italy) at 37°C for 30 min. After phenol/chloroform extraction, RNA was precipitated and the pellet dissolved in 20 µl of water. Each RNA preparation was then tested by PCR with Na_V_ primers (Table 1) to exclude the presence of residual contaminating genomic DNA. First strand cDNA was synthesized in 20 µl reaction using oligo-dt primers and ImProm-II Reverse Transcriptase (Promega, Milan, Italy) with 5 µl of RNA according to the manufacturer’s instructions. Typically, 1 µl of cDNA synthesis reaction was employed in each PCR reaction for Na_V_ amplification in taste bud preparations. Control tissues (brain, muscle and lingual epithelium) were subjected to the same procedures for RNA extraction and cDNA synthesis; however, the amount of cDNA which was employed for Na_V_ amplification was normalized to the expression level of β-actin in taste bud cDNA preparations. PCR reaction (25 μl) contained 12.5 μl of Go-Taq Master Mixes (Promega, Milan, Italy), 1 µl (25 pmoles) primers, 1 µl template cDNA and 10.5 µl water (Table 1). PCR conditions for Na_V_ cDNA amplification were: 3 min at 94°C for the initial denaturation followed by 40 cycles of 94°C for 30 sec, 54°C for 20 sec, and an extension of 72°C for 30 sec or 5 min in the last cycle. PCR conditions for β-actin cDNA amplification were: 3 min at 94°C for the initial denaturation followed by 35 cycles of 94°C for 30 sec, 62°C for 20 sec, and the extension of 72°C for 45 sec or 5 min in the last cycle. Five µl of each PCR reaction were analyzed by agarose gel electrophoresis. PCR products were quantified by densitometric analysis, processing gel images with Adobe Photoshop CS3 Extended software. All PCR products (except for β-actin) of the predicted molecular weight were extracted and purified from gel, and sequenced for validation.

**Table 1.**
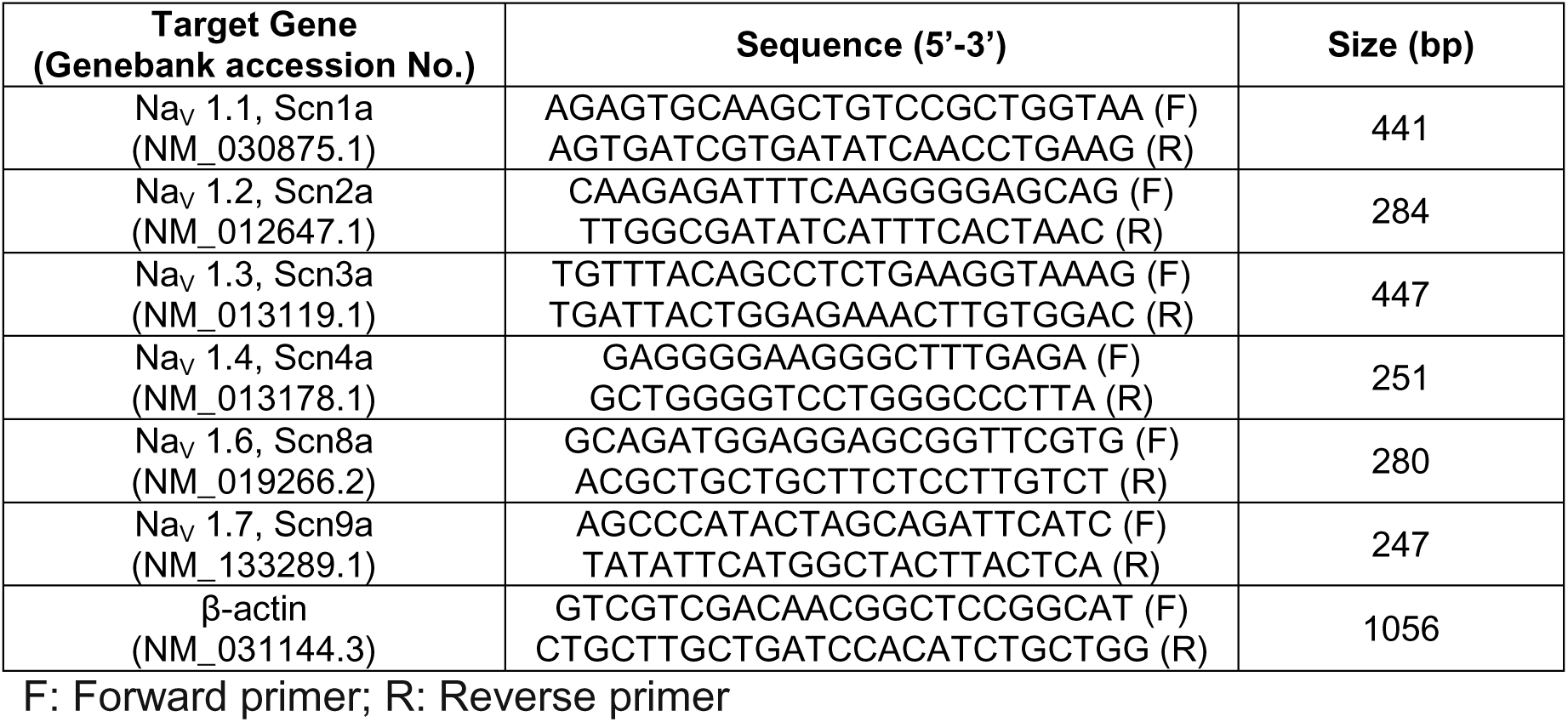
Sequences of primer sets used for RT-PCR

### Statistics

Analysis and plotting were performed using Prism 9.0 software (Graph Pad Software, La Jolla, USA). When appropriate, results were presented as medians with 95% of confidential interval (CI), unless otherwise indicated. Data comparisons were made with Mann–Whitney test. Spearman test was used to establish correlation between variables. Significance level asterisks: * = P<0.01; ** = P<0.001; *** = P<0.0001.

## Results

### Taste cells with I_Na_ and I_Am_

Taste buds in rat fungiform papillae contain distinct cell populations with specific electrophysiological properties (Bigiani & Cuoghi, 2007; Bigiani, 2015). Thus, our first goal was to identify and characterize taste cells endowed with both voltage-gated sodium channels and functional ENaCs, that is, the so-called *sodium cells* (Nomura *et al*., 2020). Using a standard voltage-clamp protocol to elicit *I_Na_* and bath application of 1 μM amiloride to monitor the amiloride response, *I_Am_*, we were able to unambiguously identify 68 sodium cells. These cells were heterogeneous with regard to the amplitude of both *I_Na_* and *I_Am_* (Fig. 1 A, B). We noticed that cells with large *I_Na_* also tended to have large *I_Am_* (Fig. 1 C). We then checked whether there was a correlation between these electrophysiological parameters. Indeed, *I_Am_* amplitude was positively correlated with *I_Na_* amplitude (Spearman correlation coefficient *r* = 0.7838, *P* < 0.0001).

**Figure 1.**
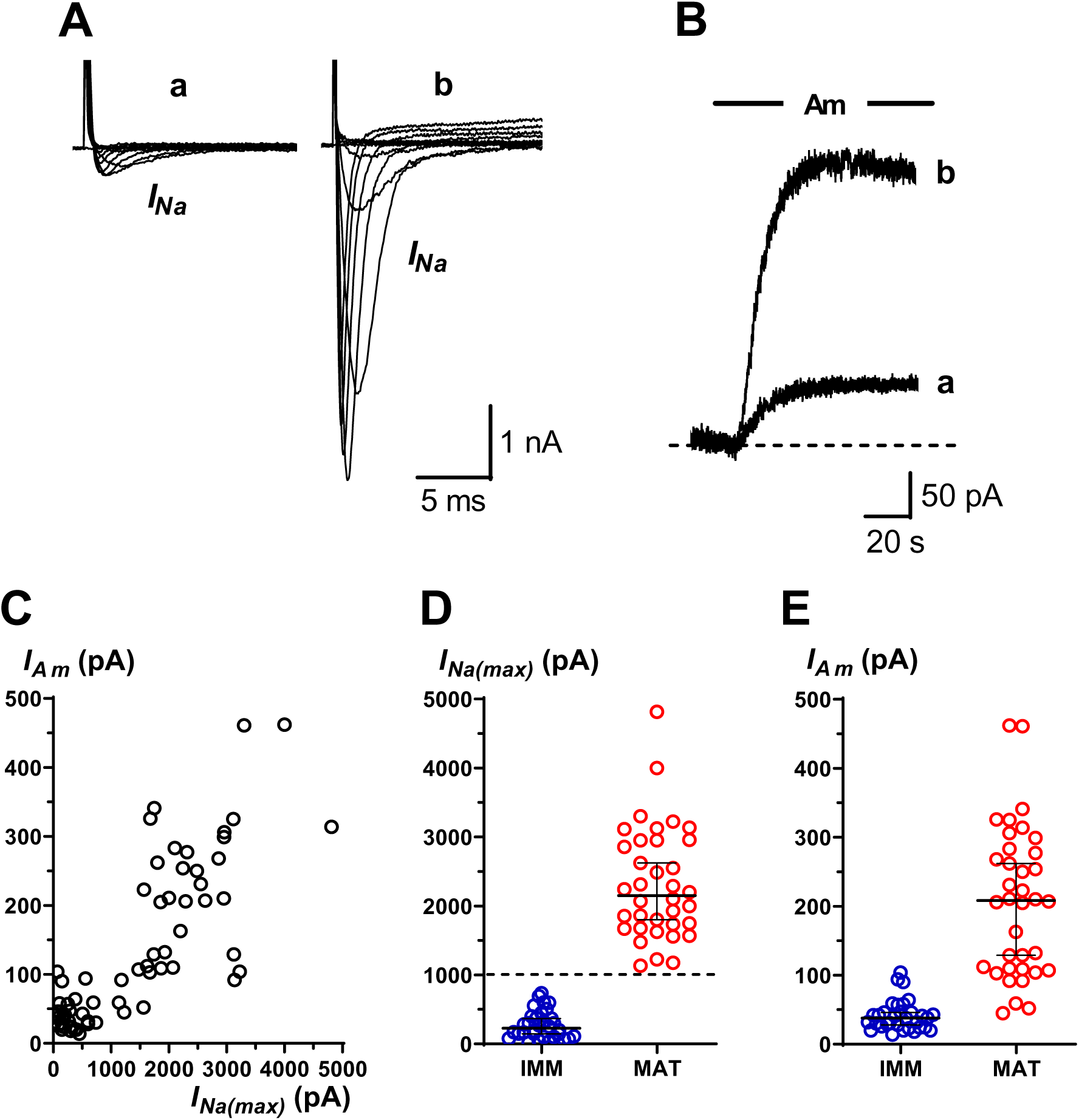
Functional cell subtyping according to the amplitude of voltage-gated sodium currents (*I_Na_*) in taste cells expressing functional ENaC. (**A**) Whole-cell, voltage-clamp recordings from a taste cell with small (a) and one with large (b) *I_Na_*. Cells were held at –80 mV and step depolarized (10-mV increments) from –70 mV to 20 mV. *I_Na_* appears as downward deflections in the current records. (**B**) For functional identification of taste cells, the effect of bath-applied 1μM amiloride (Am) was evaluated on the background membrane current when the cell was held at −80 mV (response to amiloride). These current traces were recorded from the same cells shown in A. (**C**) Scatterplot of peak amplitude of *I_Na_* vs. response to amiloride (*I_Am_*) for each recorded taste cells (n = 68). *I_Am_* was positively correlated with *I_Na_* (Spearman correlation coefficient: r = 0.7838, *P* < 0.0001). (**D**) Scatterplots of the peak values of *I_Na_* in immature (IMM) and mature (MAT) taste cells (n = 32 and 36, respectively). Dotted line indicates the current amplitude level used for subdividing cells into the two groups. Horizontal lines show medians (228 pA and 2150 pA, respectively) with 95% CI. (**E**) Scatterplots of the amplitude of the response to amiloride in immature (IMM) and mature (MAT) taste cells (n = 32 and 36, respectively). Statistical comparison with Mann–Whitney test indicated that the two distributions were significantly different (P < 0.0001). Horizontal lines show medians (38 pA and 209 pA, respectively) with 95% CI.

In the amphibian *Necturus maculosus*, *I_Na_* undergoes a gradual increase in amplitude as taste cells differentiate (Mackay-Sim *et al*., 1996). Thus, the variability of *I_Na_* in taste cells expressing ENaC was likely related to their functional maturation. Indeed, in *Necturus* taste buds, immature cells exhibit *I_Na_* with amplitude smaller than 1 nA, whereas mature cells possess *I_Na_* with amplitude larger than 1 nA (Mackay-Sim *et al*., 1996). The range of *I_Na_* amplitude in rat fungiform taste cells is similar to that found in *Necturus* (Fig. 1C) (Doolin & Gilbertson, 1996; Kossel *et al*., 1997; Bigiani & Cuoghi, 2007). Thus, we adopted the same cutoff value of 1 nA to subdivide recorded sodium cells into two groups: the first, containing cells with *I_Na_* < 1 nA, was considered “*enriched in developing cells*”, whereas the second group, containing cells with *I_Na_* > 1 nA, was considered “*enriched in fully differentiated cells*”. For simplicity, we named them as *immature* and *mature* subsets, respectively. In short, this approach allowed us to study the properties of *I_Na_* during the transition between two broad maturation states. The distributions of *I_Na_* and *I_Am_* amplitudes in these two functional subsets are shown in Fig. 1D and Fig. 1 E. Since *I_Am_* reflects the activity of ENaC, which functions as sodium receptor in rodents (Chandrashekar *et al*., 2010; Nomura *et al*., 2020), these distributions suggested that cells in the mature subset were, on average, more excitable (larger *I_Na_*) and more chemo- sensitive (larger *I_Am_*) than cells of the immature subset.

### Morphology of immature and mature taste cells

To confirm that taste cells in the two subsets were indeed at different developmental stages, we also analyzed their morphology. In histological studies, two main classes of taste cells can be identified based on their morphology (Yang *et al*., 2020; Finger & Barlow, 2021): 1) elongated spindle-shaped cells reaching the taste pore; 2) round cells in basal position or elongated cells that do not reach the taste pore (Fig. 2A). The first group contains mature taste cells, whereas the second group contains immature post-mitotic cells, which differentiate into mature elements during turnover (Barlow, 2015; Finger & Barlow, 2021). Therefore, the length of cells can provide hints about maturity, as the immature cells tend to be shorter than mature ones.

**Figure 2.**
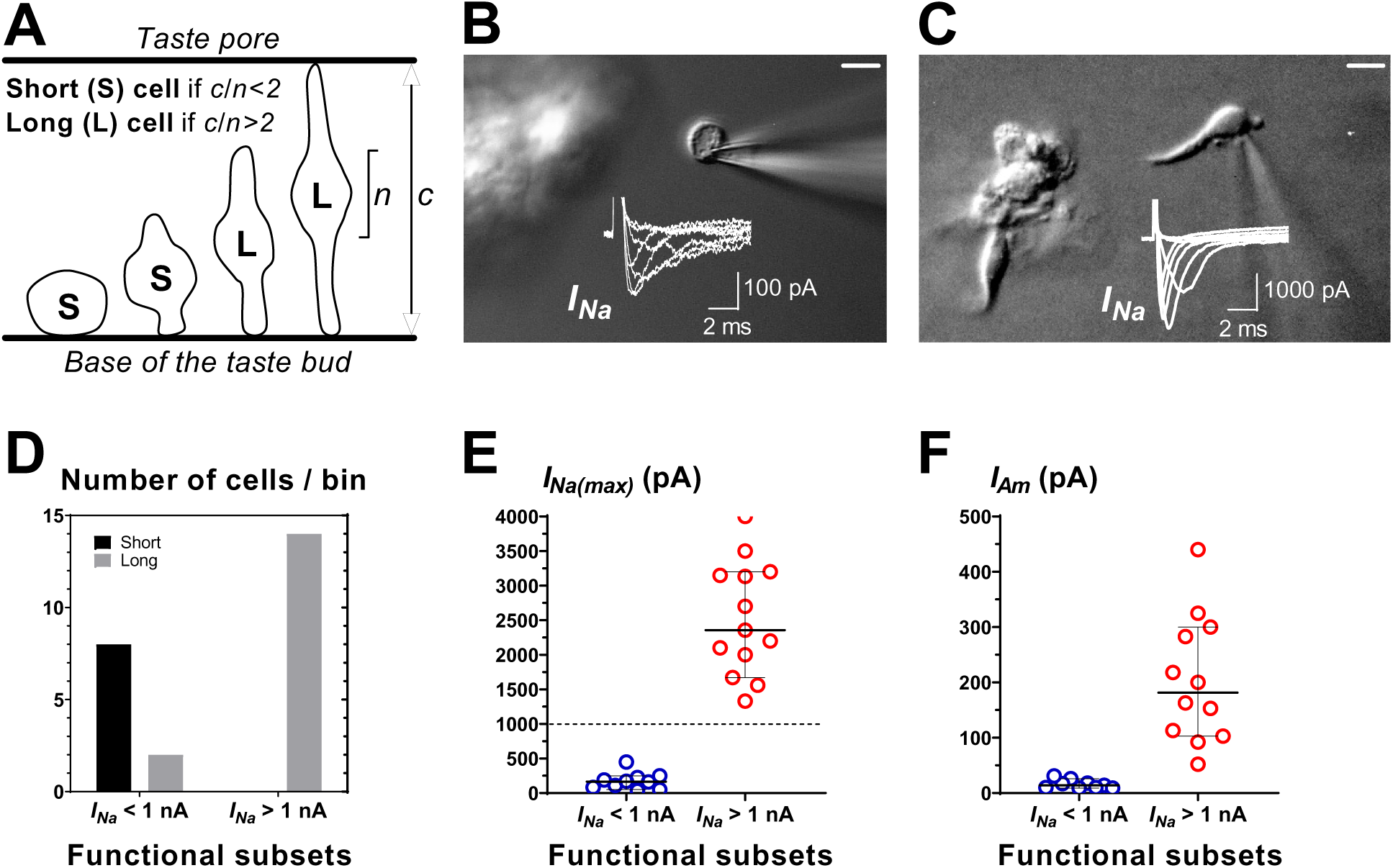
Morphology of the recorded cells. (**A**) Schematic drawing showing the change in shape and size of taste cells during turnover. By measuring *c* (total length of the cell) and *n* (length of the nuclear region), a morphometric index, *c/n*, was obtained for dividing cells into two groups. Cells with *c/n* < 2 are considered as “short” (S), cells with *c/n > 2* are considered as “long” (L). Immature cells are typically included in the “short” group, whereas mature cells belong to the “long” group. (**B**) Differential interference contrast photomicrograph depicting a “short” taste cell extracted from an isolated taste bud (left) after whole-cell recording. Note that this cell lacks any evident process. Scale bar = 5 μm. Voltage-gated sodium currents (*I_Na_*) recording from this cell is superimposed on the photomicrograph. Cell was held at −100 mV (to increase current intensity) and step depolarized (10-mV increments) from −70 mV to −20 mV. The membrane capacitance of this cell was 13.6 pF. (**C**) Differential interference contrast photomicrograph depicting a “long” taste cell extracted from an isolated taste bud (left) after whole-cell recording. Note that this cell has a long apical process. Scale bar = 5 μm. Voltage-gated sodium currents (*I_Na_*) recording from this cell is superimposed on the photomicrograph. Cell was held at −80 mV and step depolarized (10-mV increments) from –70 mV to 0 mV. The membrane capacitance of this cell was 5.2 pF. (**D**) Recorded cells were subdivided according to the *I_Na_* intensity (functional subsets) and the index *c/n* (“short” or “long”). Cells considered functionally immature (*I_Na_* < 1 nA) were mostly “short”, whereas cells considered functionally mature (*I_Na_* > 1 nA) belonged to the “long” group. (**E**) Scatterplots of the peak values of *I_Na_* in immature (*I_Na_* < 1 nA) and mature (*I_Na_* > 1 nA) taste cells identified morphologically after patch-clamp recordings. Currents were elicited from a holding potential of −80 mV. Horizontal lines show medians (165 pA and 2356 pA, respectively) with 95% CI. Dotted line indicates the current amplitude level used for subdividing cells into the two functional groups. (**F**) Scatterplots of the amplitude of the response to amiloride in immature (*I_Na_* < 1 nA) and mature (*I_Na_* < 1 nA) taste cells identified morphologically after patch-clamp recordings. Horizontal lines show medians (14 pA and 182 pA, respectively) with 95% CI. Given the low yield in obtaining identifiable taste cells after extraction from taste buds with the patch pipette, for this analysis it was not applied the rule of considering only cells with an amiloride response greater than 20 pA as for the rest of this study (see Materials and Methods). This explains the low median value for cells endowed with *I_Na_* < 1 nA.

In a separate set of recordings, we exploited these morphological features to establish the maturation stage of taste cells we recorded from. We evaluated the index “*c/n*”, where “*c*” is the total length of the cell and “*n*” is the length of the cell body (Fig. 2A). We subdivided recorded cells into two groups based on this index. Specifically, we defined a cell as “long” or “short” if *c/n* was larger or smaller than 2, respectively (Fig. 2A). This number was inferred by analyzing images of mature and immature taste cells recently described by Yang and colleagues (Yang *et al*., 2020). After recording, under direct microscope observation, we extracted the cell from the taste bud by pulling it apart gently with the patch pipette and immediately measured the *c/n* index (Fig. 2B and 2C). Only cells that maintained their original morphology after removal were considered for this analysis and doubtful cases were discarded. Although this strict criterion greatly reduced the yield in these experiments, nonetheless, we were able to uniquely assign 24 cells to one of the two morphological subgroups. The association between *I_Na_* intensity (an electrophysiological marker for cell maturity) and the *c/n* index is shown in Fig. 2D. It can be noted that cells with small *I_Na_* (n = 10) tended to be “short” (*c/n* < 2), whereas all cells with large *I_Na_* (n = 14) belonged to the “long” group (*c/n* > 2). Two immature cells (*I_Na_* < 1 nA) had a *c/n* larger than 2, that is, they were classified as “long”. This is consistent with the morphological features of developing taste cells (Yang *et al*., 2020; Finger & Barlow, 2021), which include intermediate forms (Fig. 2A). The amplitude of both *I_Na_* and *I_Am_* of identified cells is shown in Figure 2E and 2F, respectively. In summary, our electrophysiological criterion of using a cutoff value of 1 nA for *I_Na_* amplitude to distinguish two broad groups of developing taste cells was in good agreement with morphological observations for cell maturity (Yang *et al*., 2020; Finger & Barlow, 2021).

### Cell capacitance and I_Na_ density

Taste cells change significantly their morphology during development (Fig. 2A), raising the possibility that immature and mature cells might differ in cell membrane extension. This geometric dimension can be indirectly assessed by measuring the electrical capacitance of the cell, *C_M_* (Hille, 2001). Application of a voltage step to taste cells produced current transients reflecting the charging process of cell membrane. Usually, these current waveforms in immature cells were wider than in mature cells (Fig. 3A), indicating a larger *C_M_* (Fig. 3B) and therefore a larger membrane surface area. An increased *C_M_* could be due to electrical coupling between taste cells (Bigiani & Roper, 1993). In patch-clamp recordings, capacitive transient in coupled cells is characterized by two exponential decays associated with two straight lines after linearization (Bigiani & Roper, 1995). We verified whether this was the case in our conditions by linearizing the falling phase of capacitive transients (Fig. 3A, *inset*). Points could be fitted to a single straight line, ruling out the presence of additional, electrically coupled cells.

**Figure 3.**
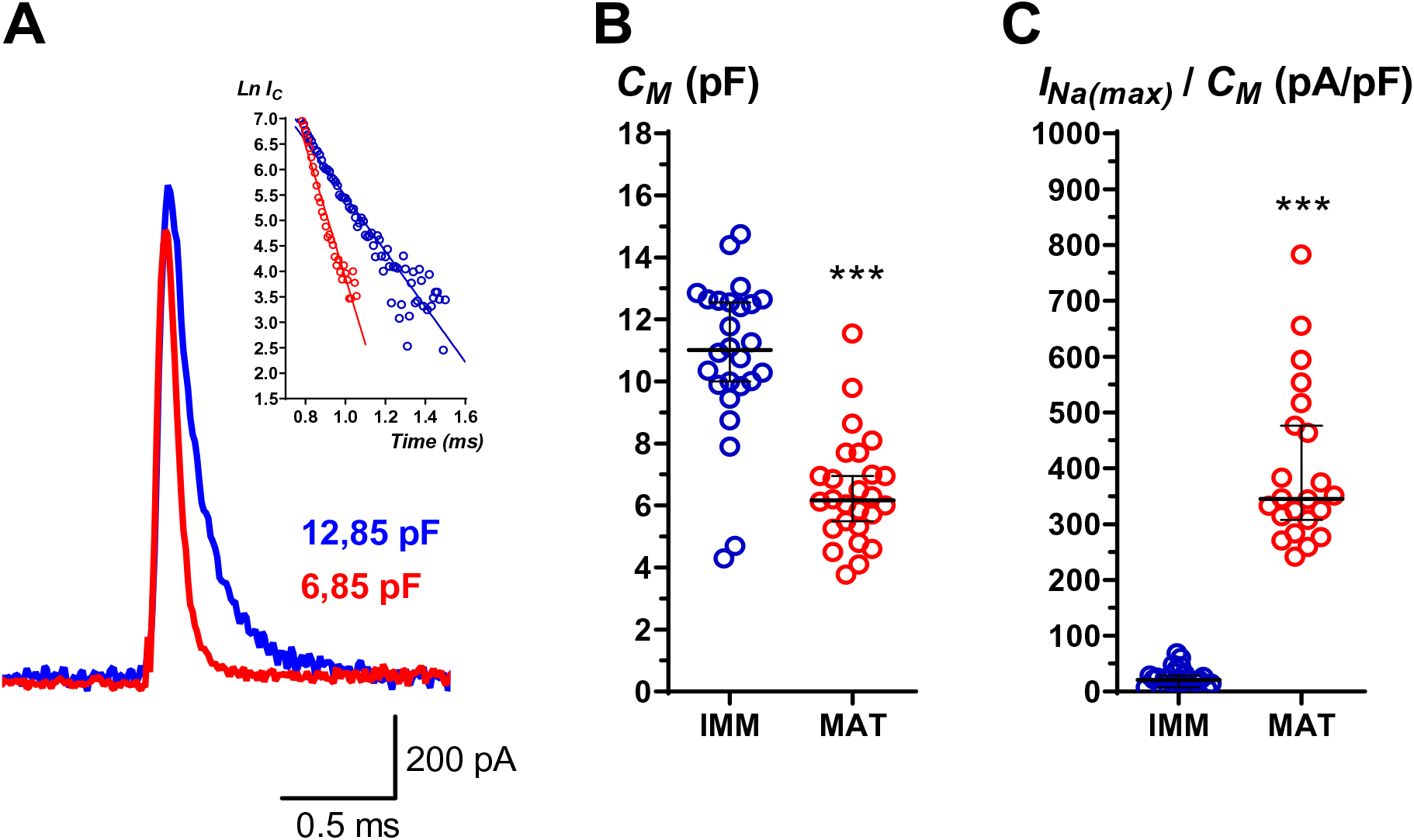
Membrane capacitance (*C_M_*) and density of voltage-gated sodium currents (*I_Na(max)_/C_M_*) in immature and mature taste cells. (**A**) Transient capacitive current elicited by a voltage step of 20 mV form a holding potential of −80 mV in an immature (blue trace) and a mature (red trace) taste cell in the presence of 1 µM amiloride. Note the slower relaxation of the current in immature cell as compared to the mature one. The access resistance was similar for the two cells, as indicated by amplitude of the transients. *Inset*: Log-plots for the relation phase of capacitive currents. Points are fitted by a single straight line, indicating that a single time constant exponential describes the relaxation phase. (**B**) Scatterplots of cell capacitance (*C_M_*) evaluated by the integral of the capacitive transients in immature (IMM) and mature (MAT) taste cells (n = 26 and 26, respectively). Horizontal lines show medians (11.0 pF and 6.2 pF, respectively) with 95% CI. Statistical comparison with Mann–Whitney test indicated that the two distributions were significantly different (P < 0.0001). (**C**) Scatterplots of the density of voltage-gated sodium current (*I_Na_/C_m_*) in immature (IMM) and mature (MAT) taste cells (n = 24 and 22, respectively). *I_Na(max)_/C_M_* is the ratio between the maximal amplitude of *I_Na_* and the value of cell capacitance (*C_M_*) for each recorded cell. Horizontal lines show medians (21 pA/pF and 345 pA/pF, respectively) with 95% CI. Statistical comparison with Mann–Whitney test indicated that the two distributions were significantly different (P < 0.0001).

*C_M_* values could obviously affect the density of *I_Na_*, that is, the ratio between *I_Na_* peak amplitude and the extension of the cell membrane defined by *C_M_* (*I_Na_/C_M_*). *I_Na_* density is an important parameter for determining the electrical excitability of the cells, as demonstrated by membrane modeling (Matzner & Devor, 1992; Platkiewicz & Brette, 2010). We found that *I_Na_/C_M_* was significantly smaller in immature cells than in mature cells (Fig. 3C). Interestingly, the change in current density appeared clear-cut, thus resembling a “quantum leap” phenomenon (Fig. 3C). In contrast, the change in current amplitude was more gradual (Fig. 1D). This finding suggests that a large *C_M_* helps keep developing cells unexcitable. It is worth noting that *I_Na_/C_M_* values are in good agreement with those found in developing neurons (Cummins *et al*., 1994; Costa, 1996; Gao & Ziskind-Conhaim, 1998; Fry, 2006) but not in *Necturus* taste cells, which exhibit an average value of ∼ 52-57 pA/pF in the mature stage (Bigiani & Roper, 1993; Mackay-Sim *et al*., 1996).

### Current-voltage (I-V) relationships

Current-voltage (*I-V*) relationships (Sontheimer & Ransom, 2002) were used to analyze the behavior of *I_Na_* as a function of the membrane potential. To this end, cells were held at −80 mV and step depolarized in 5 mV-increments (Béhé *et al*., 1990; Herness & Sun, 1995). Figure 4 (*left*) shows an example of current traces recorded from an immature (*top*) and a mature (*bottom*) cell. Current amplitude data from 16 immature cells and from 14 mature cells were then plotted against step potential to obtain the corresponding average *I-V* curves (Fig. 4, *right*). In immature cells the average peak amplitude of *I_Na_* was reached at about −20 mV, whereas in mature cells at about −10 mV. In other words, the *I-V* curve shifted towards more depolarized potentials in mature cells (Fig. 4, *right*). These findings suggested that the voltage-dependent properties of the sodium channels differed between the two cell subsets. We therefore analyzed in detail both the activation and the inactivation process.

**Figure 4.**
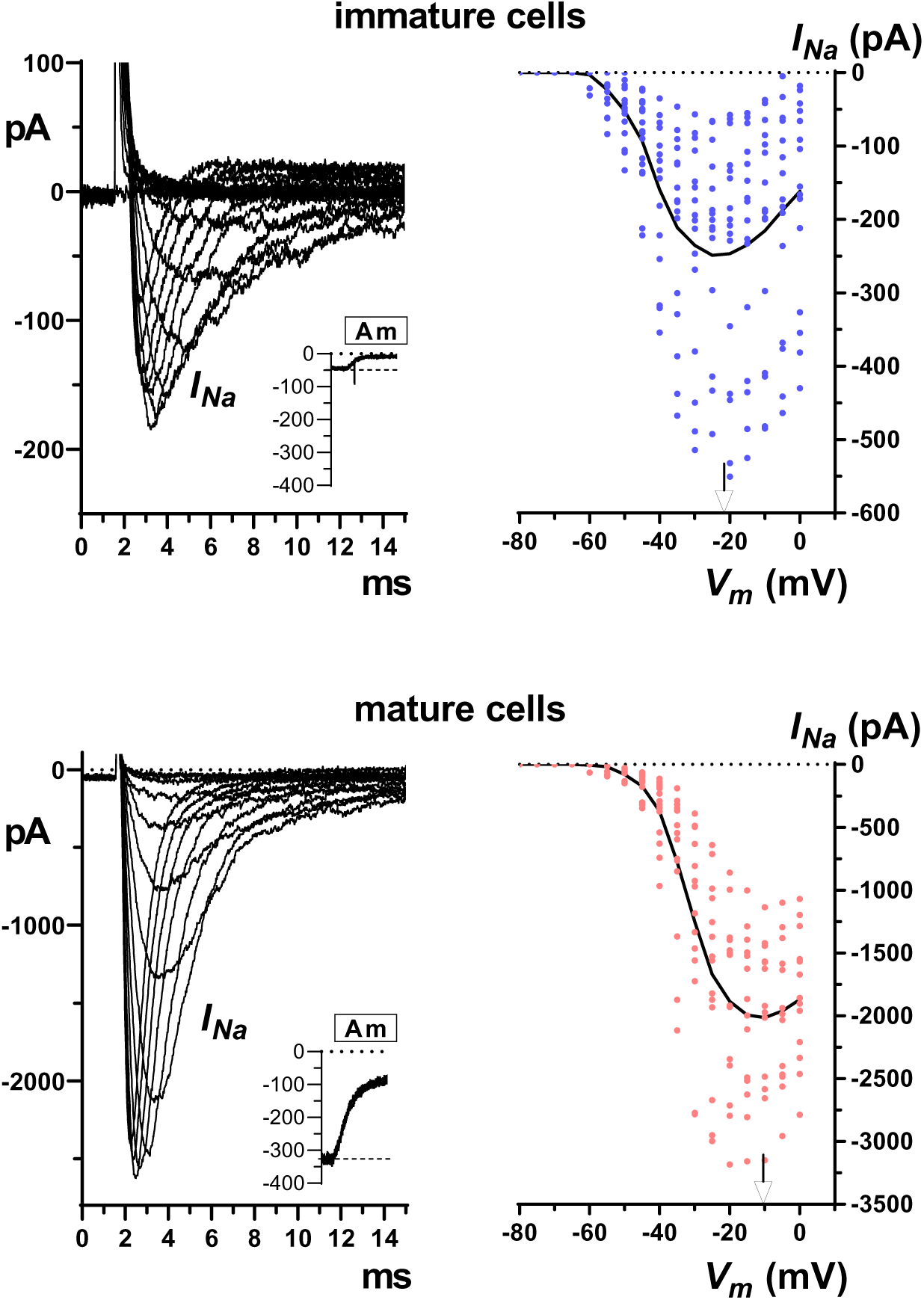
Voltage-gated sodium currents (*I_Na_*) in taste cells expressing ENaC. Whole-cell, voltage-clamp recordings from immature cells (*top*) and mature ones (*bottom*). Cells were held at −80 mV and step depolarized (5-mV increments) from −80 mV to 0 mV in the presence of 1 µM amiloride. Sample recordings are shown on the left. *I_Na_* appears as downward deflections in the current records (*left*). To obtain the corresponding current-voltage curves (*right*), the maximum amplitude of *I_Na_* for each voltage steps (*V_m_*) was plotted against the step membrane potential for 16 immature cells (blue points) and 14 mature cells (red points). Connecting lines are through mean values of the current at each membrane potential. *Arrows* point the membrane potential where *I_Na_* peaks. *Insets*: response to amiloride (Am). Y-axis: picoamperes.

### Conductance-voltage (G-V) relationships

The activation properties of *I_Na_* were evaluated by determining how membrane potential affected sodium conductance (*G_Na_*), which reflects the number of open channels. *G_Na_* was obtained by dividing the peak current amplitude at each membrane potential by the driving force for sodium ions (see Materials and Methods). Figure 5A shows sample recordings of *I_Na_* evoked at different membrane potentials (step potentials) in an immature (*top*) and in a mature (*bottom*) taste cell. Peak current amplitude was then converted to conductance, which was normalized to obtain the corresponding *G-V* plots. Data could be fitted to a Boltzmann function yielding sigmoidal curves (Fig. 5B). According to these plots, activation of voltage-gated sodium channels occurred at more positive membrane potentials (∼ 10 mV shift) in mature taste cells than in immature ones. The scatter plot for *V_0.5_* values from single cells clearly shows a significant difference in their distribution between groups (Fig. 5C). On the contrary, the *k* value in the Boltzmann relation did not change between groups (Fig. 5D).

**Figure 5.**
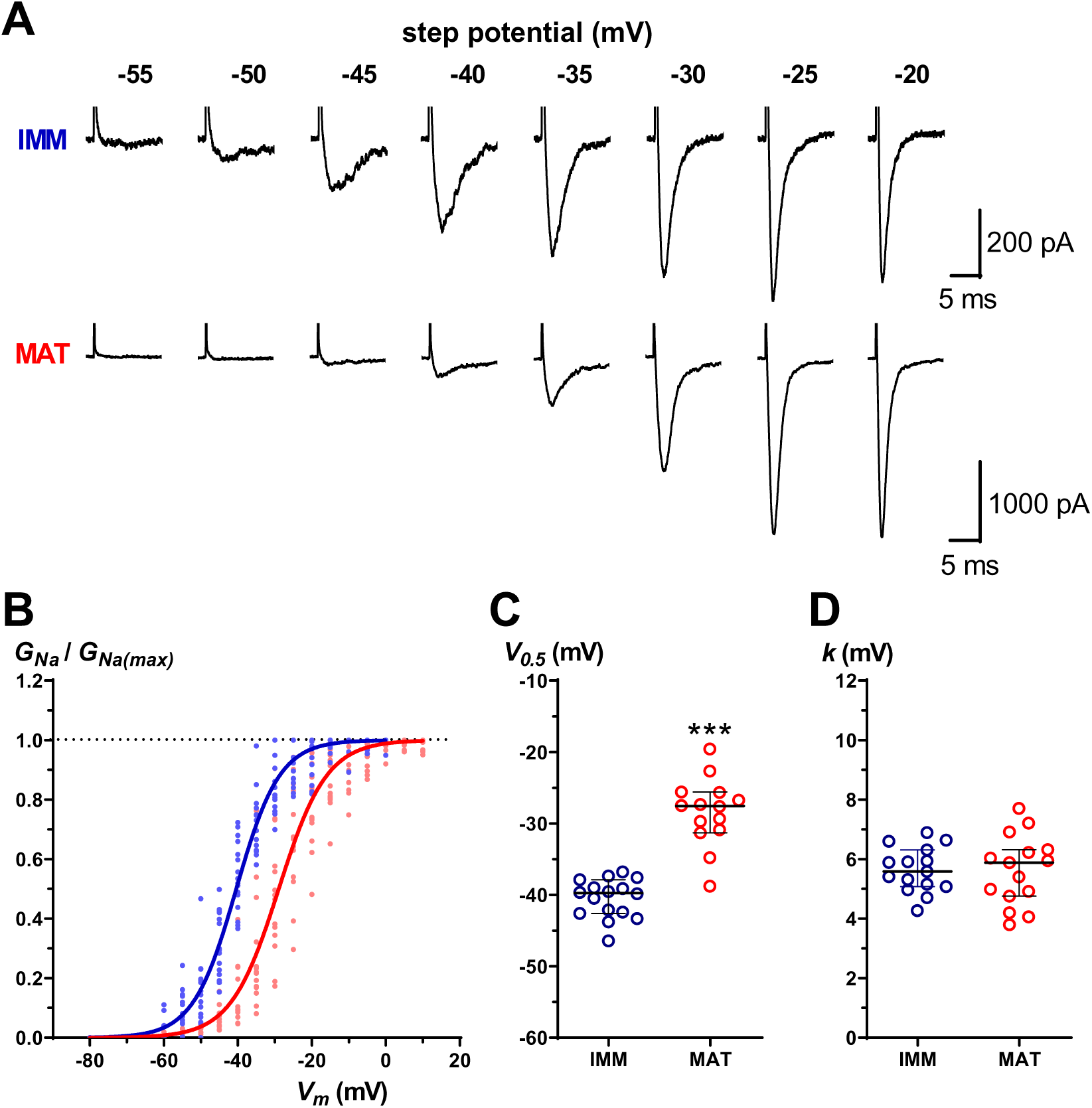
Activation of voltage-gated sodium currents (*I_Na_*) in fungiform taste cells. **A**: Sample recordings from two taste cells possessing small (*top*) and large (*bottom*) *I_Na_*. Cells were held at −80 mV and depolarized in 5-mV steps to activate sodium current in the presence of 1 µM amiloride. Step potentials are shown over recordings. **B**: Voltage-dependence of *I_Na_* activation in immature (n = 16, blue points) and mature cells (n = 14, red points). The magnitude of the current at each step potential was transformed in *G_Na_*, normalized to its maximal value, *G_Na(max)_*, and then plotted against the step potential (*V_m_*). Data were fitted to a single Boltzmann equation (R^2^: 0.9702 for immature cells and 0.9465 for mature ones). The half-maximal voltage (*V_0.5_*) was −40.4 mV and −28.9, respectively. The slope (*k*) was 5.9 mV and 6.5 mV, respectively. **C**: Scatterplots of *V_0.5_* values in taste cells endowed with small I_Na_ (< 1 nA; immature cells) and in those expressing large *I_Na_* (>1 nA; mature cells). Horizontal lines show medians with 95% CI. Median values: −39.8 mV and −27.6 mV, respectively. Statistical analysis indicated that the two distributions were significantly different (Mann–Whitney test, P < 0.0001). **D**: Scatterplots of *k* values in taste cells endowed with small *I_Na_* (< 1 nA; immature cells) and in those expressing large *I_Na_* (> 1 nA; mature cells). Horizontal lines show medians with 95% CI. Median values: 5.6 mV and 5.8 mV, respectively. Statistical analysis indicated that the two distributions were not significantly different (Mann–Whitney test, P = 0.9674).

### Steady-state inactivation

We used the standard two-pulse protocol to study *I_Na_* steady-state inactivation (*h*), which is important for the repolarizing phase of action potentials (Ulbricht, 2005). Sample recordings of *I_Na_* evoked with this stimulation protocol are shown in Fig. 6A. The ratio *I_Na(pulse)_/I_Na(max)_* was then plotted against the prepulse potential and a single Boltzmann function was used to fit the data (Fig. 6B). The voltage dependence of inactivation shifted to more positive potential values in mature cells as compared to the immature ones (∼ 10-mV shift). Furthermore, the analysis of the distribution of *V_0.5_* and *k* values revealed a significant difference between the two cell subsets (Fig. 6 C, D).

**Figure 6.**
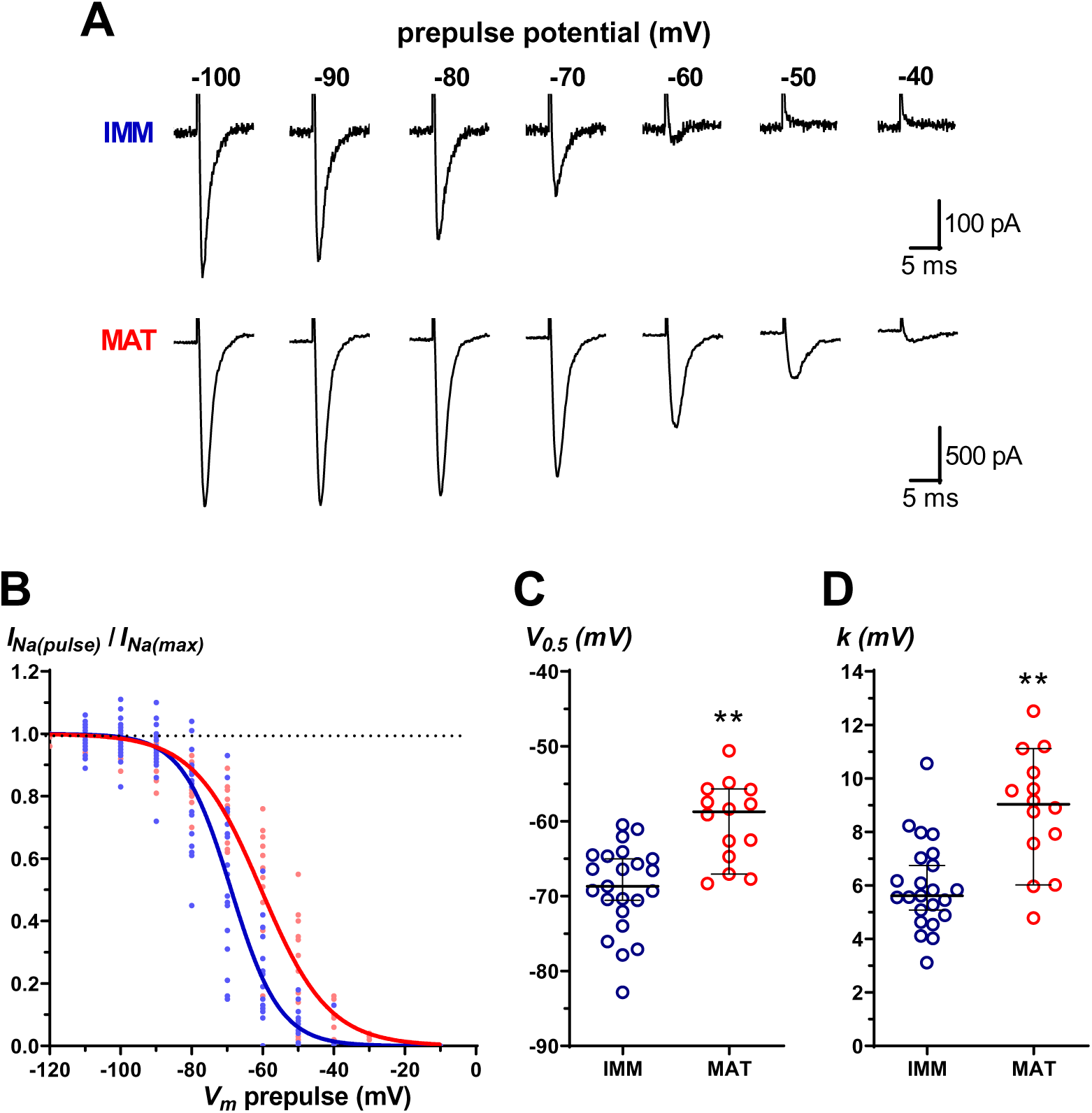
Inactivation of voltage-gated sodium currents (*I_Na_*) in fungiform taste cells. **A**: Sample recordings from two taste cells possessing small (*top*) and large (*bottom*) *I_Na_*. A standard two-pulse voltage protocol was used for this analysis. Prepulses, 500 ms in duration and of variable amplitude (from −100 mV to −20 mV), were applied prior to the test pulse to −20 mV. Cells were held at −80 mV between trials and in the presence of 1 µM amiloride. **B**: Voltage-dependence of the steady-state inactivation of I_Na_ for immature (n = 23, blue points) and mature cells (n = 14, red points). The magnitude of the current elicited by the test pulse (−20 mV) was normalized to its maximal value and plotted against the pre-pulse potential. Data were fitted to a single Boltzmann equation (R^2^: 0.9578 for immature cells, and 0.9612 for mature ones). The half-maximal voltage (V_0.5_) was −68.9 mV and −60.4 mV, respectively. The slope (k) was 6.9 mV and 9.5 mV, respectively. **C**: Scatterplots of *V_0.5_* values in taste cells endowed with small *I_Na_* (< 1 nA; immature cells) and in those expressing large *I_Na_* (> 1 nA; mature cells). Horizontal lines show median with 95% CI. Median values: −68.7 mV and −58.8 mV, respectively. Statistical analysis indicated that the two distributions were significantly different (Mann–Whitney test, P = 0.0001). **D**: Scatterplots of k values in taste cells endowed with small *I_Na_* (< 1 nA; immature cells) and in those expressing large *I_Na_* (> 1 nA; mature cells). Horizontal lines show median with 95% CI. Median values: 5.6 mV and 9.0 mV, respectively. Statistical analysis indicated that the two distributions were significantly different (Mann–Whitney test, P = 0.0002).

Unlike the activation curve, which maintained a similar slope factor (*k*) during cell maturation (Fig. 5 B and D), the inactivation curve showed a larger *k* value in mature cells than in immature cells (Fig. 6 B and D). This prompted us to further analyze the inactivation process. By studying the relationship between the *mean* values of the ratio *I_Na(pulse)_/I_Na(max)_* and the membrane potential (Fig. 7A), we found that data points from immature cells were well fitted by a single Boltzmann relationship (blue line), whereas data from mature cells were not (red line), particularly for negative potentials. Indeed, double Boltzmann expression better fitted data from mature cells (Fig. 7B, red line) as two components (“*A*” and “*B*”; see Materials and Methods) with different *V_0.5_* and *k* could be easily detected (Fig. 7B). Component “*A*” had a very hyperpolarized *V_0.5_* (*V_0.5(1)_* = −88.1 mV) and a rather large *k* (*k_(1)_* = 11.3 mV), whereas component “*B*” had a depolarized *V_0.5_* (*V_0.5(2)_* = −56.2 mV), but a *k* (*k_(2)_* = 6.8 mV) similar to that obtained with either single or double Boltzmann (6.9 mV and 6.8 mV, respectively) in immature cells. These findings suggest that the “B” component in mature cells resembled the single Boltzmann curve of the immature cells shifted by approximately 10 mV in the depolarizing direction on the voltage axis. Component “*A*”, therefore, popped up during maturation. Note that double Boltzmann fit was indistinguishable from single Boltzmann fit with data from immature cells (compare Fig. 7A with Fig. 7B). The linearization of the two datasets (immature and mature cells) allowed us to obtain a better visualization of the presence of multiple components underlying inactivation. In immature cells, a single straight line fitted the transformed data (Fig. 7C), suggesting that a single process was responsible for the steady-state inactivation in these cells. On the contrary, two different linear portions with distinct slopes could be detected for the transformed data from mature cells (Fig. 7D). This indicated that two processes were responsible for the observed steady-state inactivation in these cells.

**Figure 7.**
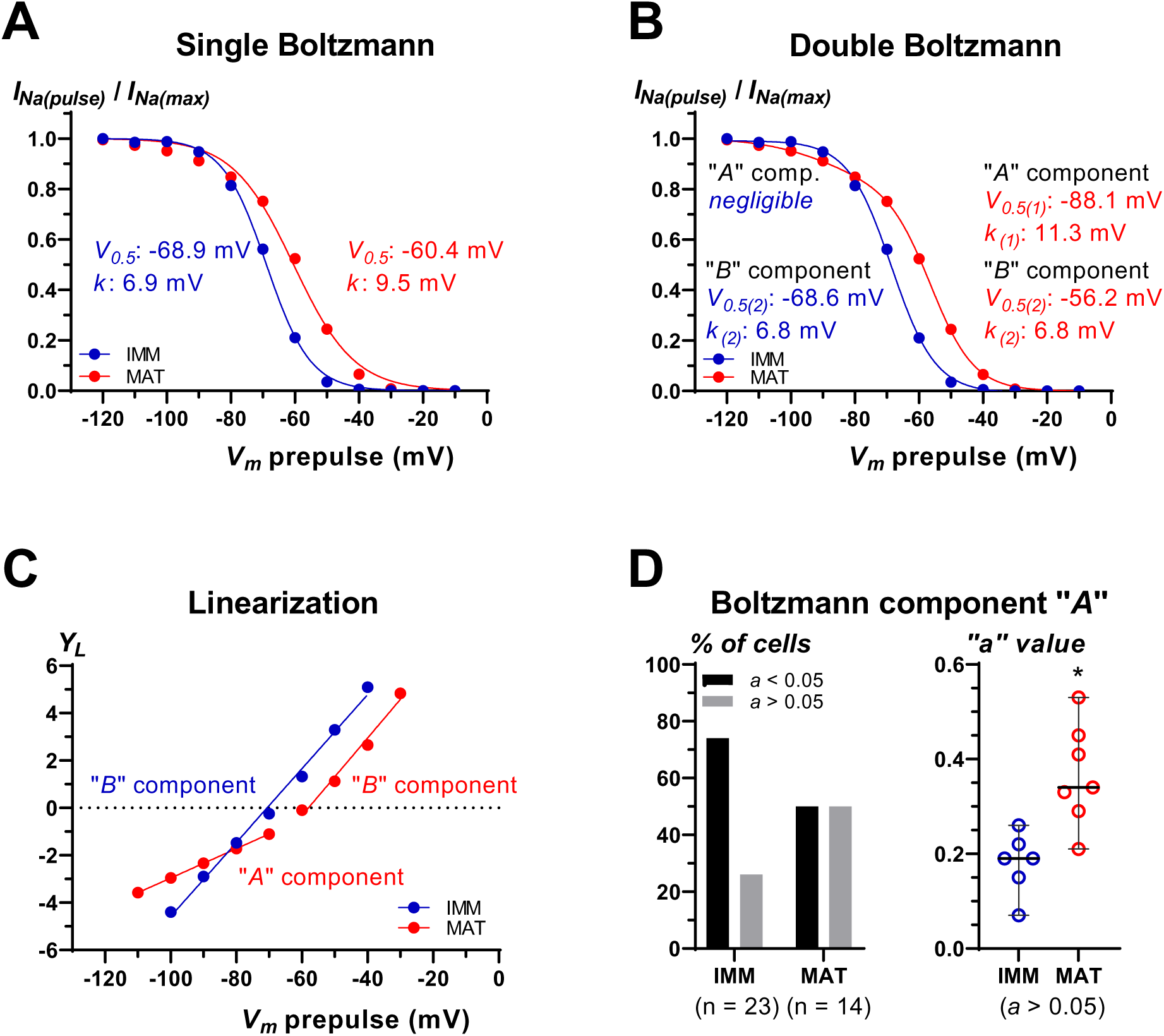
Boltzmann relationships for the steady-state inactivation of voltage-gated sodium currents (*I_Na_*) in fungiform taste cells. Points are the mean values extracted from data shown in Fig. 6B. Single Boltzmann (**A**) or double Boltzmann fits (**B**) were used to describe the shape of the curves. *I_Na_/I_Na(max)_* and *V_m_* prepulse as in Fig. 6B. Note that data from immature cells (IMM) could be fitted with a single Boltzmann equation. On the contrary, data from mature cells (MAT) were better fitted by a double Boltzmann relationship. **C**: Data linearization shows that a single straight line could fit points of immature cells. This suggested that a single process underlies inactivation process. On the contrary, data linearization shows that points of mature cells could be fitted by two straight lines. Therefore, two separate processes contributed in determining *I_Na_* inactivation in these cells. *Y’*: see Materials and Methods for details. **D**: Analysis of the component “*A*” in the double Boltzmann equation (see Materials and Methods for details). The percentage of taste cells with a distinct *A* component (*a* > 0.05) was larger in mature cells as compared to immature ones (*left*). The analysis of taste cells with *a* > 0.05 showed that the contribution of this component to the overall inactivation process was significantly higher (Mann–Whitney test, P = 0.0047) in mature cells as compared to immature ones (*right*).

Since the maturation of *I_Na_* was characterized by the appearance of a new inactivation mechanism described by the “*A*” component in the double Boltzmann function (Fig. 7B), we therefore evaluated the contribution of this component to the overall inactivation process for each single immature or mature cell. In the double Boltzmann equation, *a* may vary from 0 to 1 (see Materials and Methods). We chose a threshold value of 0.05 to establish the contribution of the new component “*A*” to the inactivation. As expected, the percentage of cells with a distinct “*A*” component (*a* > 0.05) was small in immature cells, whereas it was larger in mature cells (Fig. 7D, *left*). In addition, we found that the contribution of the “*A*” component (*a* > 0.05) to the inactivation process was less pronounced in immature cells and increased significantly in mature cells (Fig. 7D, *right*). These observations explained why fitting the average data from immature cells with a double Boltzmann function left unchanged the values of *V_0.5_* and *k* (compare blue lines in Fig. 7A and Fig. 7B). All these findings point to significant modifications in the *I_Na_* inactivation process during development.

### Voltage dependence of I_Na_ during taste cell maturation

The voltage-dependent properties of *I_Na_* are fundamental for membrane excitability (Angelino & Brenner, 2007). Our findings on activation and inactivation allowed us to gain an overall picture of how voltage dependence of *I_Na_* changed during development. Figure 8A shows the conductance curve (*G*) and inactivation curve (*h*) for both immature (*left*) and mature (*right*) cells. In both cases, it appeared to be a “window” of overlap between *G* curve and *h* curve. Additionally, the two windows were very similar, with an intersection point of *G* = *h* ≈ 0.1 in both cell subsets (Fig. 8A, dashed line). This is consistent with the value obtained in single taste cells dissociated from rat vallate/foliate papillae (Herness & Sun, 1995). Two main differences could be detected between immature and mature subsets, though. First, the intersection of *G* and *h* curves occurred at approximately −53 mV in immature cells and at approximately −43 mV in mature cells. Second, an additional component appeared in the *h* curve of mature cells (Fig. 8A, *arrow*). Clearly, these findings suggest that a change in membrane excitability occurs during maturation of taste cells in rat fungiform taste buds.

**Figure 8.**
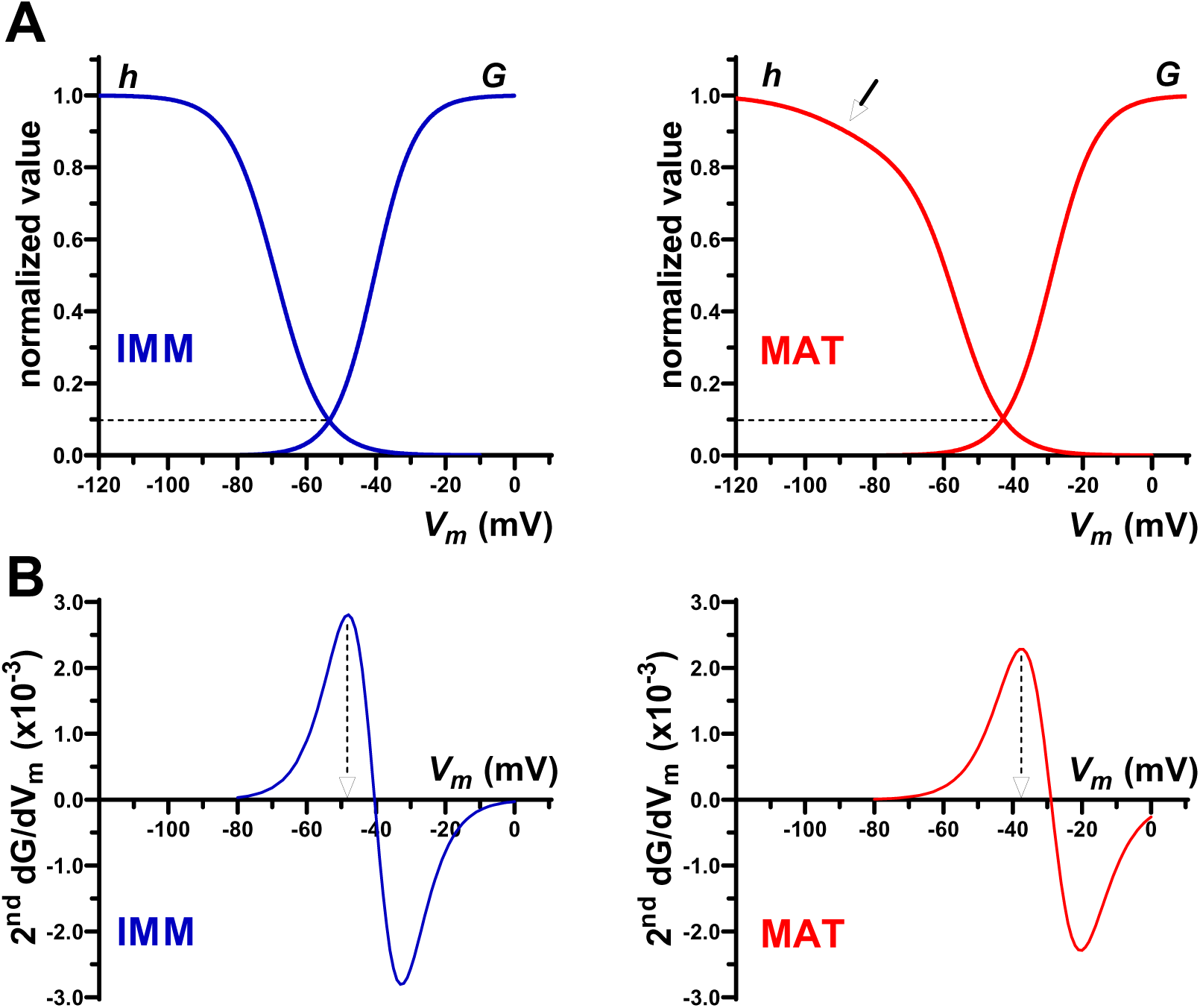
Voltage dependence of activation (*G*) and inactivation (*h*) of voltage-gated sodium currents in taste cells of the rat fungiform taste buds. **A:** A single Boltzmann relationship described the voltage-dependence of *G* in both immature (*IMM*) and mature (*MAT*) cells. Steady-state inactivation (*h*) was described by a single Boltzmann equation in immature cells (*IMM*), whereas in mature cells (*MAT*) a double Boltzmann equation was required to better fit data points. A “window” of overlap between the *G* and the *h* curves is present in both plots. Note that this window is similar in the two cell subsets, although it occurs at different membrane potentials. *Arrow*: “A” component of the double Boltzmann curve. *Dashed line*: value of the intersection point on the Y axis. **B**: Second derivative of the G functions (2^nd^ *dG/dV_m_*) shown on top. Threshold potential was defined as the membrane potential (dashed arrow) at which 2^nd^ *dG/dV_m_* had the largest positive value. Threshold potential was about −48 mV for immature cells and about −37 mV for mature cells.

The calculation of the second derivative of *G* function (2^nd^ *dG/dV_m_*; Fig. 8B) made it possible to determine the threshold potential at which the Hodgkin-Huxley cycle is triggered and opening of channels occurs with the maximal acceleration. The largest positive value of 2^nd^ *dG/dV_m_* pointed to the membrane potential at which the change rate of curve slope was greatest. As indicated by the dashed arrows in Fig. 8B, positive 2^nd^ *dG/dV_m_* peaked at about −48 mV for immature cells and about −37 mV for mature cells. This suggests that the threshold for action potential firing shift towards more positive membrane potentials during cell maturation.

### Persistent (non-inactivating) sodium current

The analysis of the voltage dependence of inactivation indicated that *I_Na_* in taste cells could be due to multicomponent combinations of sodium channels. Consistent with this hypothesis, we noticed that current decay differed somehow between immature and mature cells. We then compare the inactivation kinetics at −30 mV (Raman *et al*., 1997). Figure 9A shows an example of the time course of the current inactivation in the two functional subsets. While current decayed to zero in immature cells, current in mature cells was characterized by a slow and late component resembling the persistent current described in other excitable cells (Raman *et al*., 1997; Smith *et al*., 1998; Rush *et al*., 2005). The amplitude of this persistent current was small compared to *I_Na_* peak amplitude; nevertheless, it could be detected in most mature taste cells (13 out of 16; Fig. 9B red circles), whereas was essentially absent in immature cells (1 out 16 cells; Fig. 9B blue circles). Therefore, it is conceivable that this component of the sodium current is added during development. Although a detailed analysis of the kinetics of current inactivation was beyond the scope of this study, however, we found that current decay could be approximately fitted by a single exponential function in both immature and mature cells (Fig. 9C). The corresponding time constant did not differ significantly between subsets (Fig. 9D). This finding suggests that the fast phase of the current decay (first two milliseconds delimited by dashed lines in Fig. 9C, where current traces are approximately linear and corresponding to about a 37% decrease) is likely due to sodium channels with similar properties in both immature and mature cells.

**Figure 9.**
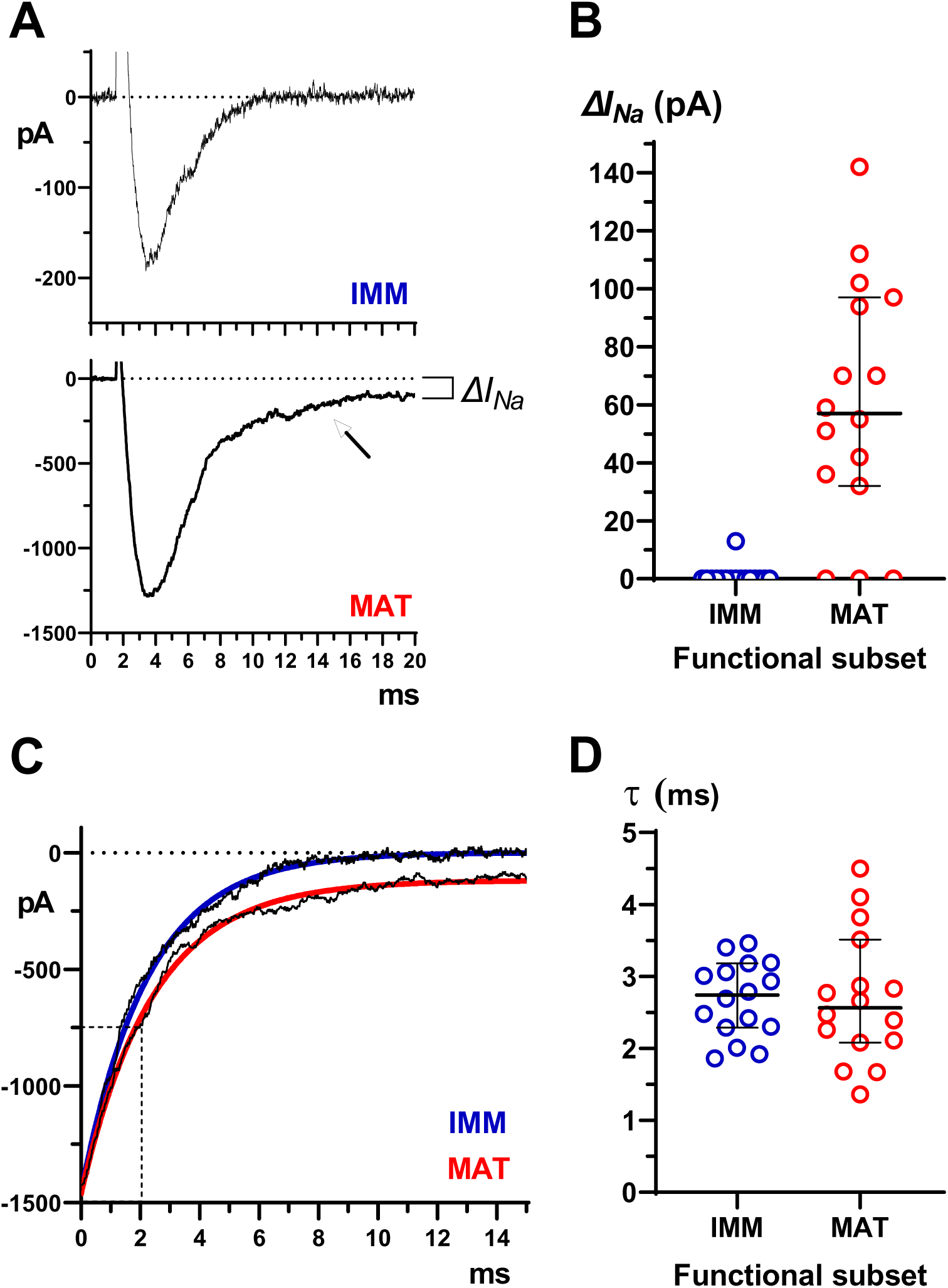
Persistent *I_Na_* in mature taste cells. **A**: Currents evoked by step depolarization to −30 mV (holding potential −80 mV) in an immature (IMM) and a mature (MAT) taste cell. Note that the current decays more slowly in the mature cell (*arrow*). Δ*I_Na_*: amplitude of late (persistent) current measured at 20 ms from the beginning of step depolarization. **B**: Scatterplot of Δ*I_Na_* values in immature (IMM; n = 16) and mature (MAT: n = 16) taste cells. Horizontal lines show medians with 95% CI. For immature cells, median is hidden by data points due to the low occurrence of persistent current in these cells (see text for details). **C**: A single exponential in both immature (IMM) and mature (MAT) cells could fit current decay due to inactivation. Current trace from immature cell was scaled to the amplitude of the current from mature cell. The dashed lines delimit approximately the initial rapid decay of *I_Na_*. **D**. Scatterplots of time constant (*τ*) values for the exponential function describing the current decay due to inactivation in immature and mature cells (n = 16 and 16, respectively). Horizontal lines show median with 95% CI. Median values: 2.74 ms mV and 2.57 ms, respectively. Statistical analysis indicated that the two distributions were not significantly different (Mann–Whitney test, P = 0.6963).

### Na_V_ α-subunits expressed in fungiform taste buds

Voltage-gated sodium channels (Na_V_) consist of an α-subunit, which provides the conductive pore, and one or more auxiliary β-subunits (Catterall *et al*., 2005). The nine isoforms of Na_V_ α-subunits (1.1 – 1.9) exhibit distinctive biophysical properties, such as the voltage dependence of inactivation, when heterologously expressed in mammalian cell lines. Na_V_1.6 and Na_V_1.7 analyzed by a two-pulse protocol with 500-ms prepulses exhibit fairly negative *V_0.5_* value for inactivation (−74 mV and −78 mV, respectively) which might be compatible with the *V_0.5_* of second Boltzmann component in the inactivation curve that appears in mature fungiform taste cells (Fig. 7B) (Cummins *et al*., 1998; Zhao *et al*., 2011). Interestingly, Na_V_1.6 is also responsible for the persistent currents described in other excitable cells (O’Brien & Meisler, 2013), a feature that we observed in mature taste cells (Fig. 9). Moreover, the *V_0.5_* value for inactivation that we found in immature fungiform taste cells (−68.6 mV; Fig. 7B) was in the range of those reported for recombinant Na_V_1.3 and Na_V_1.4 expressed in mammalian cell lines (−65/-72 mV) (Cummins *et al*., 1998; Cummins *et al*., 2001; Vilin *et al*., 2012). Therefore, we seek to identity the Na_V_ α-subunit isoforms expressed in fungiform taste buds through mRNA detection. In rodents, chemo-sensitive taste cells exclusively possess TTX-sensitive Na_V_ channels (Béhé *et al*., 1990; Herness & Sun, 1995; Furue & Yoshii, 1997; Gao *et al*., 2009; Kimura *et al*., 2014; Ohtubo, 2021), therefore, we restricted the analysis to the six TTX-sensitive Na_V_ α-subunits, namely Na_V_1.1, Na_V_1.2, Na_V_1.3, Na_V_1.4, Na_V_1.6, and Na_V_1.7 (Catterall *et al*., 2005). For this assay, mRNA was extracted from ten pools of fungiform papillae preparations, each of which contained isolated taste buds from three animals. RT-PCR was then employed to identify the α-subunit isoforms expressed in each pool. Importantly, since this approach provided information on the mRNA from the whole taste buds, it is likely that Na_V_ mRNAs from (taste) cells not expressing ENaC were also revealed.

Figure 10 shows the results of the RT-PCR analysis. Four α-subunits were detected in our tissue preparations (Fig. 10A), albeit with different occurrences and relative abundance. The most frequently expressed subunits were Na_V_1.3, Na_V_1.4, and Na_V_1.7, whereas Na_V_1.6 occurred less frequently (Fig. 10B). Furthermore, Na_V_1.3, Na_V_1.4, and Na_V_1.7 appeared to be expressed more abundantly than Na_V_1.6 when compared to the expression of the same Na_V_ isoforms in brain as a control (Fig. 10B). In contrast, Na_V_1.1 and Na_V_1.2 were never detected in any pool (n = 10; Fig. 10 B).

**Figure 10.**
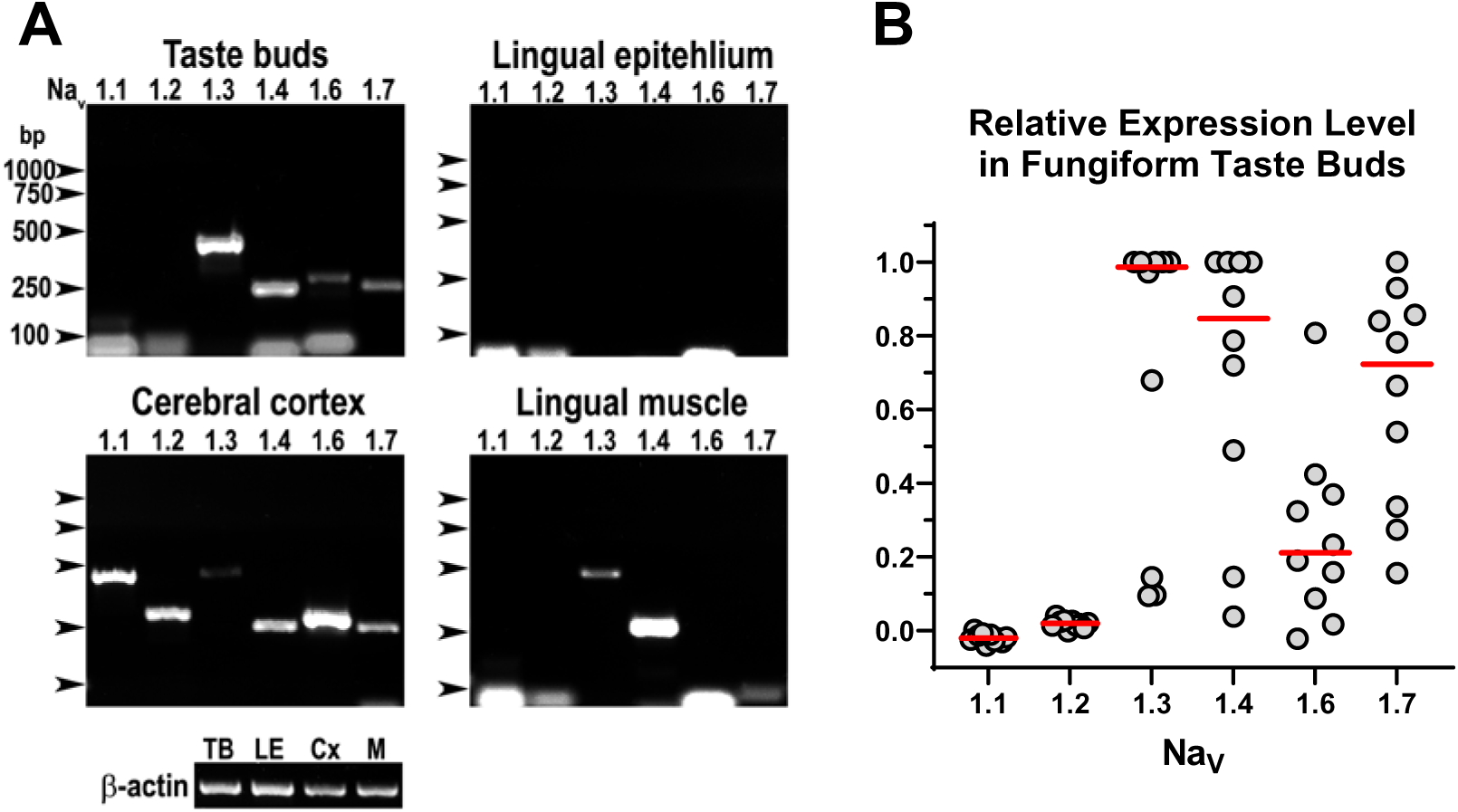
RT-PCR expression analysis of the Na_V_ isoforms in rat fungiform taste buds. **A:** Electrophoresis of RT-PCR of total RNA extracted from fungiform taste buds of 3 rats. Shown is the expression of the Na_V_ isoforms (1.1, 1.2, 1.3, 1.4, 1.6, 1.7). Negative and positive controls for Na_V_ isoform expression were from lingual non sensory epithelium, cerebral cortex, and lingual skeletal muscle. The amount of reverse transcribed RNAs was normalized to β–actin expression level before amplification with Na_V_ primers. Arrows indicate the position of the DNA markers; bp, base pair **B:** Densitometric analysis of Na_V_ expression in RNA from isolated taste buds. Shown are the relative expression levels of the Na_V_ isoforms in each preparation (n = 10). Circles with values of 1 refer to the most intense electrophoretic band in each preparation. Red bars represent the median values of relative expression for each Na_V_ isoforms.

## Discussion

In this study we analyzed the biophysical properties of *I_Na_* in a functional cell line of rat fungiform taste buds, namely, the so-called sodium cells (Nomura *et al*., 2020). These electrically excitable cells express the amiloride-sensitive, epithelial sodium channel (ENaC), a sodium receptor for detecting low-medium salt concentrations in rodents (Chandrashekar *et al*., 2010; Nomura *et al*., 2020). Our goal was to study the *I_Na_* modifications during cell turnover, which occurs continuously in taste buds (Barlow, 2015). Due to this dynamic process, sodium cells were electrophysiologically and morphologically heterogeneous (Fig. 1). Therefore, we simplified the analysis by subdividing recorded cells into two groups according to *I_Na_* amplitude, a hallmark of cell maturity as demonstrated in *Necturus* taste cells (Mackay-Sim *et al*., 1996) and in many other excitable tissues, *e.g.* (Cummins *et al*., 1994; Costa, 1996; Gao & Ziskind-Conhaim, 1998; Cummins *et al*., 2002; Donnelly, 2011; Calvigioni *et al*., 2017). Accordingly, the group containing cells with *I_Na_* < 1 nA was considered “*enriched in developing (immature) cells*”, whereas the group containing cells with *I_Na_* > 1 nA was considered “*enriched in fully differentiated (mature) cells*”. Histological studies have shown that mature taste cells are spindle-shaped elements extending approximatively from the base to the apical end of taste buds, where they make contact with chemical stimuli. Immature taste cells, on the contrary, are either ovoid or elongated cells that do not reach the apical end of the taste buds. Thus, morphological maturation of taste cells consists in switching from a rounded shape to a spindle-like shape, with intermediate forms (Feng *et al*., 2014; Barlow, 2015; Yang *et al*., 2020). As demonstrated by analyzing the shape of recorded cells (Fig. 2), grouping of taste cells by *I_Na_* amplitude supported the validity of our approach. Clearly, this represents an oversimplification of the actual situation occurring in the taste buds; nevertheless, our functional criterion for cell grouping proved very useful in providing meaningful information on the transition from immature to mature state. To our knowledge, this is the first study describing functional changes in mammalian taste cells during turnover.

### C_M_ in the transition “immature ➔ mature”

The changes in taste cell shape during turnover raise an obvious question: does the membrane surface area change in parallel with the morphological variation? The patch-clamp recording technique allows at evaluating *C_M_*, which is directly related to the surface area of the cell membrane (Hille, 2001). In *Necturus* taste buds, *C_M_* increases as cell differentiation proceeds due to an increase in the size of taste cells (Mackay-Sim *et al*., 1996). An increase in *C_M_* during development seems to be a general rule also in many types of neurons and it is associated to an increase in cell size, *e.g.* (Gao & Ziskind-Conhaim, 1998; Cummins *et al*., 2002; Picken Bahrey & Moody, 2003; Rimmer & Harper, 2006; Donnelly, 2011; Ogawa *et al*., 2017). Unexpectedly, in rat taste buds, we found a larger *C_M_* in immature cells than in mature cells although they shared a seemingly similar cell body (Fig. 2 B,C).

A decreased *C_M_* in mature taste cells could result from a reduction in spatial charging capability during application of a voltage step. Indeed, this is expected if patched cell exhibits a complex morphology, with cellular branches and processes (Golowasch *et al*., 2009). Although mature taste cells are spindle-shaped, they are small and the charging process of the cell membrane during a voltage step can be described by a single exponential function (Fig. 3A), suggesting that under our recording conditions taste cells were electrotonically compact. A recent paper shows that immature, post-mitotic taste cells have an uneven and irregular membrane, whereas mature chemo-sensitive taste cells possess a regular and smooth membrane (Yang *et al*., 2020). Thus, it is conceivable that a larger *C_M_* in immature taste cells reflects a particular organization of their cell membrane, which is wider due to the irregular morphology. The observed *C_M_* change from immature to mature state suggests reorganization of cell membrane in parallel with morphological changes of the cell shape.

Due to cell turnover, taste buds contain differentiating (immature) cells, differentiated (mature) cells, but also degenerating cells (Feng *et al*., 2014). Could it be possible that some immature cells were actually dying elements? Although we cannot entirely rule out this possibility, however, cell death is reportedly accompanied by a reduction in membrane capacitance (Mărgineanu & Van Driessche, 1990; Kalauzi *et al*., 2015). Thus, the larger membrane capacitance in immature cells (Fig. 3B) does not support the above prediction.

### I_Na_ in the transition “immature ➔ mature”

The ability to generate action potentials is a key functional property of chemo-sensitive taste cells, including sodium cells, as it allows them to release the neurotransmitter onto nerve fibers (Romanov *et al*., 2012; Nomura *et al*., 2020; Kolesnikov, 2021). *I_Na_* is responsible for the rapid depolarizing upstroke characterizing the action potentials in excitable taste cells (Béhé *et al*., 1990; Chen *et al*., 1996; Bigiani *et al*., 2002; Ma *et al*., 2017; Ohtubo, 2021). Our findings provide evidence for a *functional transition* from immature to mature state involving changes in *I_Na_* density (defined as the ratio between *I_Na_* peak amplitude and *C_M_*), and in the voltage dependence of activation and inactivation.

In excitable cells, including the *Necturus* taste cells, as *I_Na_* increases, *C_M_* either increases or does not change during maturation (Cummins *et al*., 1994; Mackay-Sim *et al*., 1996; Gao & Ziskind-Conhaim, 1998; Cummins *et al*., 2002; Picken Bahrey & Moody, 2003; Donnelly, 2011). Indeed, in rat taste cells, *I_Na_* significantly increased during transition from immature to mature cells; however, *C_M_* unexpectedly decreased. Therefore, *C_M_* determined a clear separation between the two cell subsets in terms of *I_Na_* density resembling a “quantum leap” (Fig. 3C). Of course, this could be the consequence of pooling cells into only two developmental groups (Costa, 1996; Gao & Ziskind-Conhaim, 1998). If more developmental stages are considered, then current density increases more smoothly (Cummins *et al*., 1994; Fry, 2006). Nonetheless, the strong reduction of *C_M_* in taste cells could suggest that increasing *I_Na_* amplitude alone is not sufficient to ensure adequate electrical excitability in mature taste cells as it is for neurons.

In excitable cells, *I_Na_* maturation is characterized by changes in the voltage dependence of activation and inactivation (Cummins *et al*., 1994; Costa, 1996; Gao & Ziskind-Conhaim, 1998; Fry, 2006; Donnelly, 2011; Yu *et al*., 2011; Tadros *et al*., 2015; Calvigioni *et al*., 2017). It is believed that these changes are necessary to adapt membrane excitability to the functional properties of mature cells. We found that also in sodium cells, the transition from immature to mature state was accompanied by changes in activation and inactivation properties. Since *I_Na_* is produced by the activity of a population of ion channels, it is possible to study the biophysical behavior of *I_Na_* in terms of a two-state Boltzmann distribution. The Boltzmann relationship allows one to obtain key parameters, *V_0.5_* and *k*, related to the voltage at which half of the channels are open and to the number of gating charges involved in the operation of the channels, respectively (Pusch, 2006). In other words, *V_0.5_* and *k* provide information on the voltage-dependent properties of sodium channels (Angelino & Brenner, 2007).

*I_Na_* activation showed significant differences between immature and mature taste cells. According to *V_0.5_*, there was a ∼10-mV depolarizing shift in the activation curve during cell maturation (Fig. 5C), whereas *k* did not change (Fig. 5D). Since data could be fitted by a single Boltzmann equation (Fig. 5B), it is reasonable to assume that in both cell subsets activation was due to a simple voltage-dependent mechanism that switches ion channels from closed state to open state. *I_Na_* maturation was therefore characterized by a change in voltage sensitivity (*V_0.5_*) but not in the number of gating charges responsible for channel opening (*k*) in the activation process. In other excitable cells, the activation curve typically undergoes a ∼10-mV hyperpolarizing voltage shift during development, *e.g.* (Cummins *et al*., 1994; Costa, 1996; Fry, 2006; Donnelly, 2011). Thus, our finding of a depolarizing shift was quite unexpected. In terms of membrane excitability, this means that during maturation the threshold for action potential generation moved to more depolarized potentials (Fig. 5B and 8); however, this depolarizing shift was fully consistent with the measured threshold potential for action potential generation in electrically excitable taste cells. For example, in both rat and mouse taste cells from the vallate/foliate papillae, the membrane potential requires a value of about −40 mV / −35 mV to elicit an action potential (Chen *et al*., 1996; Bigiani *et al*., 2002; Romanov *et al*., 2012; Ma *et al*., 2017). A similar value for action potential threshold is observed also in mouse fungiform sodium cells during stimulation with Na^+^ (Nomura *et al*., 2020). Our estimation of the membrane potential at which 2^nd^ *dG/dV_m_* had a positive peak (about −37 mV; Fig. 8B) was therefore in good agreement with the above measurements. On the other hand, immature cells that appear more excitable are likely unable of producing action potentials due to the very low density of *I_Na_* (Fig. 3C). Again, *C_M_* seems to be a key factor in determining membrane excitability in differentiating taste cells.

Unlike *V_0.5_*, the slope factor *k* in the activation curve did not change during the transition from immature to mature state (Fig. 5D). The values we estimated (about 6 mV; Fig. 5D) were in good agreement with those observed in taste cells from the vallate/foliate papillae of the rat (Herness & Sun, 1995) or the mouse (Ma *et al*., 2017; Ohtubo, 2021). Also in developing neurons, *k* does not change and has a value of about 5-8 mV (Cummins *et al*., 1994; Costa, 1996; Fry, 2006; Donnelly, 2011). *k* is inversely related to the “apparent gating valence”, which provides information on the number of gating charges in the protein responsible for switching the channel from closed to open configuration upon depolarization (Pusch, 2006). It is tempting to speculate that during maturation, the sodium channels responsible for *I_Na_* did not change their molecular features involved in channel opening, but rather their sensitivity to the membrane voltage, as indicated by *V_0.5_*.

Inactivation is an important process during action potential repolarization. It depends on a “ball-and-chain” mechanism involving the operation of a cytoplasmic domain of the Na^+^-channel α-subunit that occludes the conducting central pore (Ulbricht, 2005; Savio-Galimberti *et al*., 2012). Like the activation process, also *I_Na_* inactivation properties changed during the transition from immature to mature state. Boltzmann analysis of the data obtained with a standard two-pulse voltage protocol showed that the inactivation curve underwent a ∼10-mV shift in the depolarizing direction (Fig. 6 B). This finding was quite unexpected as in developing neurons either a 7-11 mV hyperpolarizing shift (Cummins *et al*., 1994; Costa, 1996) or no changes at all (Cummins *et al*., 2002; Fry, 2006; Donnelly, 2011) have been documented. Interestingly, also the slope factor (*k*) changed significantly during the maturation of taste cells, from a median value of 5.6 mV in immature cells to a median value of 9.0 mV in mature cells (Fig. 6D). Other studies on rat taste cells have reported average *k* values of 6.5 mV (Béhé *et al*., 1990) and 7.3 mV (Herness & Sun, 1995) whereas, in developing neurons, *k* does not change during maturation and exhibits an average value of 6-7 mV (Cummins *et al*., 1994; Costa, 1996; Fry, 2006) or 10-11 mV (Donnelly, 2011). All these studies use a single Boltzmann relation to describe the voltage dependence of inactivation. Interestingly, we found that in mature sodium cells of rat fungiform papillae, the inactivation curve was more adequately described by a double Boltzmann relation (Fig. 7B). One component (*“A”*) was characterized by a hyperpolarized *V_0.5(1)_* of about −88 mV and a large *k_(1)_* of about 11 mV (Fig. 7B). This component was negligible in immature cells (Fig. 7B). The second component (*“B”*) was characterized by a depolarized *V_0.5(2)_* of about −56 mV, and a *k_(2)_* of about 7 mV similar to that evaluated for immature cells (Fig. 7B). These findings suggest that during the transition from immature to mature state, a significant reorganization of *I_Na_* inactivation does occur, with the appearance of a new component (*“A”*) and possibly the depolarizing shift of the component already present in immature cells (*“B”*).

### Na_V_ α-subunits

According to the RT-PCR analysis, different Na_V_ α-subunits are reportedly expressed in mouse taste buds from fungiform papillae, namely Na_V_1.2, Na_V_1.3, Na_V_1.5, Na_V_1.6 and Na_V_1.7 (Gao *et al*., 2009; Ohtubo, 2021). In rat fungiform taste buds, however, the expression pattern of the Na_V_ isoforms appears to be somewhat different. In fact, whereas Na_V_1.3, Na_V_1.6 and Na_V_1.7 are significantly expressed, Na_V_1.2 could not be detected. Additionally, rat taste bud cells also express Na_V_1.4. This latter finding was unexpected as Na_V_1.4 is skeletal muscle-specific (Catterall *et al*., 2005). Although differences between mouse and rat are reported for signaling molecules (Yoshida *et al*., 2009b), however, we cannot explain the expression of this isoform in rat taste buds. It is plausible that Na_V_1.4 mRNA originates from contamination with myocytes of the lingual muscle that may have occurred during taste bud isolation; however, we were unable to detect significant Na_V_1.4 expression levels in the control lingual epithelium that underwent the same treatment.

As reported above, a double Boltzmann function provided a better fit for *I_Na_* inactivation in mature taste cells (Fig. 7B). Since Na_V_ α-subunits may differ for the voltage dependence of inactivation (Catterall *et al*., 2005), the double Boltzmann could be interpreted as due to the presence of two distinct sodium channels with *V_0.5_* of −88 mV and −56 mV, respectively. However, *V_0.5_* of inactivation may vary considerably across studies according to the duration of the pre-pulse in the two-pulse protocol. Long pre-pulses tend to shift the inactivation curve towards more negative membrane potentials (Chen *et al*., 2000; Meadows *et al*., 2002). If we limit the analysis to published data obtained by using 500-ms prepulse (our experimental conditions) and mammalian cell lines heterologously expressing Na_V_ α-subunits, then we can make the following considerations. First, none of the expressed recombinant α-subunits exhibits a *V_0.5_* for inactivation consistent with either the “*A*” component or the “*B*” component described by the double Boltzmann fitting for mature taste cells (Fig. 7B). Second, *V_0.5_* for inactivation in immature cells (−68.6 mV; Fig. 7B) is in the range of the *V_0.5_* found for Na_V_1.3 and Na_V_1.4 expressed in mammalian cell lines, namely −65 mV / −72 mV (Cummins *et al*., 1998; Cummins *et al*., 2001; Vilin *et al*., 2012). Therefore, the expression of a new Na_V_ α-subunit that may affect the overall inactivation process does not seem a reasonable explanation. Variations in the expression of Na_V_ β-subunits can greatly affect the functioning of voltage-gated sodium channels (Patino & Isom, 2010; Savio-Galimberti *et al*., 2012). Indeed, these regulatory proteins produce a depolarizing shift of inactivation as well as activation (Qu *et al*., 2001; Shah *et al*., 2001; Chen *et al*., 2002). Interestingly, the co-expression of α- and β-subunits may determine the appearance of a second component in the inactivation curve, which requires fitting with double Boltzmann equation (Shah *et al*., 2001). Since the expression of β-subunits is developmentally regulated (Patino & Isom, 2010), it is tempting to speculate that transition from immature to mature state in taste cells may coincide with the expression of these subunits. Future studies are needed to verify this possibility.

Our data confirm the expression of Na_V_1.6 in fungiform taste buds, which is responsible for the persistent sodium current in other systems (O’Brien & Meisler, 2013). This molecular evidence is consistent with the observation of a second, slow component in the inactivation kinetics of sodium current in mature taste cells (Fig. 9).

### Functional implications

Overall, our findings indicate that the voltage dependence of both activation and inactivation shifted in the depolarizing direction during maturation of sodium cells (Fig. 8). Does this change affect the membrane excitability of sodium cells? In our experiments, we applied voltage-clamp techniques to analyze the behavior of a membrane current, *I_Na_*. However, the membrane potential of taste cells *in situ* is unclamped and the ability to fire action potentials depends both on the voltage dependence of sodium channel properties and on an adequate and stable resting potential (Angelino & Brenner, 2007). Cell resting potential is a key factor for membrane excitability because it determines the fraction of sodium channels available to be opened by membrane depolarization. The experimental conditions in our recordings (Cs^+^ substituting K^+^ in the pipette solution; extracellular amiloride to block ENaC) were not suitable for determining this parameter. In taste cells without functional ENaC, the resting potential measured as zero-current potential (*V_0_*) in patch-clamp recordings ranges from −35 mV to −65 mV (Béhé *et al*., 1990; Chen *et al*., 1996; Furue & Yoshii, 1997; Bigiani *et al*., 2002; Medler *et al*., 2003; Noguchi *et al*., 2003; Ma *et al*., 2017). However, these measurements likely provide underestimates, due to the limitations of the patch-clamp technique when dealing with small cells (Barry & Lynch, 1991). Indeed, current injection to hold the membrane at about −70/-80 mV is often required for eliciting action potential with depolarizing current pulses, *e.g.* (Chen *et al*., 1996; Bigiani *et al*., 2002; Ma *et al*., 2017). This suggests that resting potential should have quite negative values in electrically excitable taste cells, including *sodium cells* (Nomura *et al*., 2020). Hence, the voltage dependence of *I_Na_* activation and inactivation described here are fully compatible with a membrane able to fire action potentials with a threshold of about −40 mV, the point in the activation curve where *G_Na_* starts to increase quickly (Fig. 8B, *right*). It is more difficult to predict membrane excitability in immature taste cells, as no direct information on their resting potential is available. In a study on the postnatal development of mouse taste cells (Bigiani *et al*., 2002), it was found that *V_0_* was significantly depolarized (about 20 mV) in taste cells from juvenile mice (age: 4-7 days) compared to adults (age: > 28 days). This suggests that the resting potential becomes more negative during development and that immature taste cells were probably endowed with a more depolarized resting potential than mature cells. Clearly, this would further limit the ability to fire action potentials because the fraction of sodium channels in the inactivated state would be quite large (Fig. 8). Furthermore, the large *C_M_* of immature cells, keeping the *I_Na_* density extremely low, could represent a development strategy to prevent the manifestation of electrical excitability before maturation is complete.

If during maturation the resting potential of taste cells becomes more negative, why do activation and inactivation curves shift towards more depolarized potentials? According to our data (Fig. 8A), it is clear that the interrelationship between activation and inactivation is already determined earlier in the maturation, as the “window” of overlap between the two curves is similar in both immature and mature cells. It is possible that the depolarized shift is related to the functioning of mature taste cells endowed with ENaCs. Since these channels are constitutively open, it is likely that the sodium content of the saliva may produce a stationary inward current (Gilbertson & Zhang, 1998). In turn, this ENaC-mediated current may cause a sustained membrane depolarization that adapts the cell to salivary sodium. Shifting both activation and inactivation curves toward more depolarized potentials may allow the adapted taste cell to fire action potentials when the sodium concentration in the saliva increases (Avenet & Lindemann, 1991). This is a critical aspect of the functioning of mature sodium cells, which need action potentials to trigger the voltage-dependent, non-vesicular release of the neurotransmitter, ATP (Nomura *et al*., 2020). In this regard, it is worth noting that the presence of a persistent sodium current (Na_V_1.6) may be important in defining the pattern of action potential discharge once the threshold is reached. Indeed, Na_V_1.6 sodium current plays a key role in regulating firing in other excitable cells (O’Brien & Meisler, 2013). In Purkinje neurons, for example, subthreshold sodium current mediated by Na_V_1.6 channels allows repetitive firing (Raman *et al*., 1997). Mathematical modeling of ATP release has shown that repetitive firing is required for optimal secretion in taste cells (Kolesnikov, 2021). Thus, Na_V_1.6 may represent an important current component to boost electrical signaling in mature sodium cells.

Taste cells sensing sweet, bitter, and umami compounds (collectively indicated as Type II cells) use action potentials to release the neurotransmitter, ATP, onto nerve endings (Taruno *et al*., 2013). Thus, an obvious question is whether changes in the biophysical properties of *I_Na_* described for sodium cells occur also in these cells. It is worth nothing that both sodium cells and Type II cells require the same transcription factor *Skn-1a* for their generation, suggesting that sodium cells may represent a subset of Type II cells (Ohmoto *et al*., 2020). It is therefore tempting to speculate that perhaps also electrical excitability in canonical Type II cells might undergo maturation process similar to those described here for sodium cells during turnover.

## Conclusions

Voltage-gated sodium current in taste cells with functional ENaC (sodium cells) undergoes complex changes during turnover, which include an increase in current density, a depolarizing shift in the voltage dependence of activation and inactivation, and the appearance of new current components. These changes are consistent with the ability of mature taste cells to generate action potentials for sustaining synaptic communication with afferent nerve endings.

As a final note, we would like to emphasize that the voltage dependence of *I_Na_*, while important, does not represent the only *I_Na_* feature that determines the membrane behavior in response to stimulus-induced depolarization. Also the kinetics properties of activation and inactivation are critical to establish sodium channel functioning and its impact on firing characteristics (Angelino & Brenner, 2007). We have only briefly touched upon this important aspect of the biophysics of *I_Na_* in taste cells. Nonetheless, this study represents a starting point for improving our understanding of how membrane excitability changes during turnover in mammalian taste cells.

## Acknowledgments

We are grateful to Mr. Claudio Frigeri for his excellent technical assistance. This work was supported in part by Università di Modena e Reggio Emilia (FAR Dipartimentale 2018 and 2019).

## Author contributions

AB conceived this study, designed and performed the electrophysiological experiments; RT designed and performed the molecular biology experiments; AB, RT, LB analyzed data and wrote the paper.

